# Efflux pumps mediate changes to fundamental bacterial physiology via membrane potential

**DOI:** 10.1101/2023.04.03.535035

**Authors:** Emily E Whittle, Oluwatosin Orababa, Alexander Osgerby, Sarah J Element, Jessica MA Blair, Tim W Overton

## Abstract

Efflux pumps are well known to be an important mechanism for removing noxious substances such as antibiotics from bacteria. Given that many antibiotics function by accumulating inside bacteria, efflux pumps contribute to resistance. Efflux pump inactivation is a potential strategy to combat antimicrobial resistance, as bacteria would not be able to pump out antibiotics. We recently discovered that the impact of loss of efflux function is only apparent in actively growing cells. We demonstrated that the global transcriptome of *Salmonella* Typhimurium is drastically altered during slower growth leading to stationary phase cells having a re-modelled, less permeable, envelope that prevents antibiotics entering the cell. Here, we investigated the effects of deleting the major efflux pump of *Salmonella* Typhimurium, AcrB, on global gene transcription across growth. We revealed that an *acrB* knockout entered stationary phase later than the wild type strain SL1344, and displayed increased and prolonged expression of genes responsible for anaerobic energy metabolism. We devised a model linking efflux and membrane potential, whereby deactivation of AcrB prevents influx of protons across the inner membrane and thereby hyperpolarisation. Knockout or deactivation of AcrB was demonstrated to increase membrane potential. We propose that the global transcription regulator ArcBA senses changes to the redox state of the quinol pool (linked to the membrane potential of the bacterium) and coordinates the shift from exponential to stationary phase via the key master regulators RpoS, Rsd, and Rmf. Inactivation of efflux pumps therefore influences the fundamental physiology of *Salmonella*, with likely impacts on multiple phenotypes.

**Importance:** We demonstrate for the first time that deactivation of efflux pumps brings about changes to gross bacterial physiology and metabolism. Rather than simply being a response to noxious substances, efflux pumps appear to play a key role in maintenance of membrane potential and thereby energy metabolism. This discovery suggests that efflux pump inhibition or inactivation might have unforeseen positive consequences on antibiotic activity. Given that stationary phase bacteria are more resistant to antibiotic uptake, late entry into stationary phase would prolong antibiotic accumulation by bacteria. Furthermore, membrane hyperpolarisation could result in increased generation of reactive species proposed to be important for the activity of some antibiotics. Finally, changes in gross physiology could also explain the decreased virulence of efflux mutants.

## Introduction

Many antibiotics function by accumulating inside bacteria and inhibiting essential metabolic functions such as protein synthesis, DNA repair, or cell wall synthesis (1). Bacteria can therefore protect themselves against these antibiotics by decreasing intracellular accumulation. There are two main mechanisms for this: removal of antibiotics from inside the cell via efflux pumps (2); and prevention of influx of antibiotics across the envelope (3). The major efflux pump in *S*. Typhimurium is AcrAB-TolC, a member of the RND family (4) comprising the transporter AcrB, the periplasmic adaptor protein AcrA, and the outer membrane channel TolC. AcrAB-TolC has been shown to have a wide substrate range and mediate resistance to multiple antibiotics. Loss of AcrB function through mutation or inhibition has been shown to make *S*. Typhimurium hyper-susceptible to antibiotics as well as avirulent, unable to form biofilm and less motile (5, 6); major changes to the physiology of the organism.

In an attempt to understand the physiological consequences of loss of efflux function previous studies have compared global gene expression in the *S*. Typhimurium wild type and efflux mutants (5–7). Wang-Kan *et al*. compared an *acrB* D408A mutant (defective in AcrB efflux function) with the wild type *S*. Typhimurium SL1344, both in exponential phase growth in MOPS minimal medium (6) and stationary phase growth in rich LB broth (7). The latter study also compared gene expression in *E. coli* K-12 MG1655 and its *acrB* D408A mutant, and characterised the exometabolome and endometabolome of each of the four strains in order to better understand the substrates of the AcrAB-TolC pump. The effect of deletion of *acrD*, encoding an alternative efflux pump with narrower substrate specificity to AcrB, on global gene expression has also been characterised (8). However, each of these studies compared gene expression at a single timepoint.

In a previous study (9) we investigated the dynamic balance between efflux and prevention of influx of antibiotics across the Gram-negative bacteria and revealed that efflux is more important during rapid exponential phase growth as the envelope is more fluid and antibiotics can pass into the bacterium more readily. However, in stationary phase, the envelope is remodelled and becomes less amenable to antibiotic entry, thus preventing intracellular accumulation, meaning that efflux has less impact. We used RNA-Seq in *Salmonella* Typhimurium SL1344 to map global gene expression as bacteria transitioned from exponential to stationary phase, highlighting gene expression changes potentially responsible for remodelling multiple layers of the envelope. Work by others has also revealed that prevention of influx and active efflux have different relative contributions during antimicrobial resistance development across Gram-negative pathogens (10, 11).

Given that the relative impact of efflux varies during bacterial growth, in the present study, we tracked gene expression at 1, 3 and 5 hours growth in the wild type and an Δ*acrB* mutant to enable the dynamics of gene expression to be compared. We sought to better understand the physiological changes brought about by deletion of *acrB*, and to differentiate between the direct effects of AcrB as an efflux pump (eg removal of compounds from the bacterium) and the secondary effects brought about by the deletion.

We show that deletion of *acrB* has wide-ranging effects on physiology and we propose and test a model that links efflux, membrane potential, redox state of the cell, and entry into stationary phase. This suggests that efflux pumps are deeply embedded in bacterial physiology, and loss of AcrB function perturbs the transition from exponential phase to stationary phase physiology. We also demonstrate the benefits of transcriptome analysis across growth, using samples taken at multiple timepoints, to discover novel aspects of gene regulation and physiology.

## Results

We previously published the results of a transcriptome analysis showing transition of the wild type *S*. Typhimurium strain SL1344 from exponential to stationary phase (9). Samples were taken after 1, 3, and 5 hours growth in MOPS minimal medium, and gene expression was tracked using RNA-Seq across growth. This study revealed changes to aspects of physiology as bacteria transitioned from the rapid growth of exponential phase to the slower growth of stationary phase. We specifically focused on changes in gene expression responsible for remodelling the bacterial envelope from a relatively permeable structure in rapid growth to a far less permeable state in stationary phase. This decreased envelope permeability is likely a key contributor to why stationary phase bacteria are more resistant to multiple antibiotics (12, 13) as well as other stressors (14, 15).

In the present work, we expanded this study by repeating these transcriptomics experiments in an *S*. Typhimurium Δ*acrB* mutant (9), similarly taking samples after 1, 3, and 5 hours growth in MOPS minimal medium. The two strains grew similarly under these conditions. First, we compared how changes in gene expression over time differed between the wild type and Δ*acrB* strains. Gene expression was compared between 1 h and 3 h, 1 h and 5 h, and 3 h and 5 h timepoints for each strain (Figure 1). This revealed that more genes were differentially regulated early in growth (the 1 h vs 3 h comparison) in the wild type than in the Δ*acrB* mutant. Comparing later timepoints, there are more differentially expressed genes for the 3 h vs 5 h comparison in the Δ*acrB* mutant than the wild type. This suggests that changes in gene expression occur later in the Δ*acrB* mutant than the wild type.

**Figure 1.**
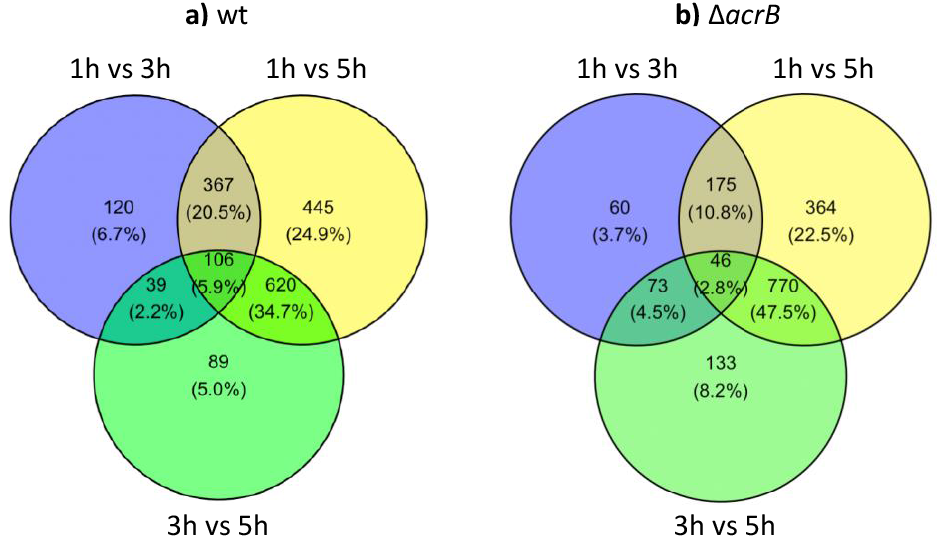
Venn diagrams comparing gene expression in the (a) wild type and (b) Δ*acrB* strain. Number of genes differentially expressed (log_2_ foldchange ≥ 1.5 and p_adj_ < 0.05) for each comparison is shown.

We wanted to understand why expression of more genes changed between 1 h and 3 h in the wild type than in the Δ*acrB* strain, and the effect this would have on physiology. We classified genes into three groups (Figure 2) based on their log_2_ fold change (log_2_FC) from 1 h to 3 h in each strain, and their Δlog_2_FC value, calculated by subtracting the Δ*acrB* log_2_FC from the wild type log_2_FC. A positive Δlog_2_FC value denotes a gene that is more upregulated or less downregulated in the Δ*acrB* strain than the wild type. Genes with a negative Δlog_2_FC value are more upregulated or less downregulated in the wild type.

**Figure 2.**
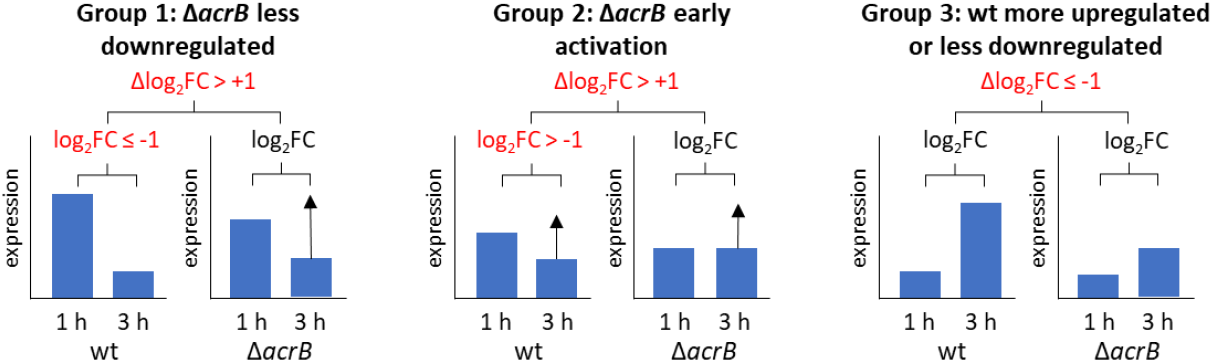
Data analysis strategy to characterise differentially expressed genes. For each gene, transcript levels at 1 h and 3 h were compared and a log_2_ fold-change (log_2_FC) value calculated. The log_2_ fold-change for the Δ*acrB* mutant was subtracted from the wt log_2_FC to give a Δlog_2_FCvalue indicating the difference in regulation between the two strains. A positive Δlog_2_ value indicates a gene is more upregulated or less downregulated in Δ*acrB* for the 1 h vs 3 h timepoints. A negative Δlog_2_ value indicates that a gene is more upregulated or less downregulated in the wild type.

Group 1 comprises genes that were downregulated from 1 h to 3 h in the WT (log_2_FC ≤ -1) and less downregulated in the Δ*acrB* strain, thereby having a Δlog_2_FC of ≥ 1. Group 2 contains genes that were not downregulated from 1 h to 3 h in the wt (log_2_FC > -1) but had a Δlog_2_FC of ≥ 1, thereby were more upregulated or less downregulated in the Δ*acrB* mutant. Group 3 contains genes that were more upregulated from 1 h to 3 h in the wild type than in the Δ*acrB* mutant. Genes with a p_adj_ > 0.05 (non-significant) for the 1 h to 3 h timepoints in either wt or Δ*acrB* were omitted from this analysis.

### Group one – late repression in the ΔacrB strain

The 166 genes in group one are more downregulated between 1 h and 3 h in the wild type than the Δ*acrB* strain (Supplemental Table S1). The largest categories of genes in this group are those encoding tRNAs (34 genes) and ribosomal proteins and other proteins involved in translation (33 genes). Many of these genes (including the tRNA genes) are known to be downregulated during the entry into stationary phase by the stringent response, mediated by ppGpp (16). For many of these genes, expression is downregulated in the wild type from 1 h to 3 h and then does not change significantly from 3 h to 5 h, whereas in the Δ*acrB* strain expression does not change from 1 h to 3 h but decreases from 3 h to 5 h. This suggests that the Δ*acrB* strain enters stationary phase later than the wild type SL1344 strain.

Fifteen genes in this group are involved in envelope functions including inner membrane protein insertion, outer membrane lipoprotein trafficking, LPS and lipid A synthesis, and peptidoglycan synthesis and remodelling. The peptidoglycan-related genes *ldtB, mrcA* and *pbpG* are implicated in exponential phase functions (9, 17). Five genes involved in nucleotide metabolism and 6 genes in amino acid metabolism, also important during rapid growth, are also in this group, reinforcing the idea that stationary phase transition is delayed in the Δ*acrB* strain.

Four genes in group one are related to polyamines, specifically putrescine import and spermidine synthesis. Recently, the efflux regulator AcrR (which represses expression of *acrAB*) was shown to bind to three polyamines and regulate aspects of polyamine efflux and metabolism in *E. coli* (18). Seven genes of *Salmonella* pathogenicity island 1 (SPI-1), encoding a type III secretion system required for invasion of epithelial cells, are in group one. The SPI-1 regulatory network is very complex (19) and comprises transcription factors encoded both within and outside SPI-1. Expression of SPI-1 is bistable (20) and is influenced by multiple environmental stimuli including osmolarity, oxygen tension, bile, [Mg^2+^], and pH. Two genes in SPI-11, implicated in survival in the macrophage (21), are also in group one. Alteration in SPI expression would likely influence virulence, which is known to be decreased upon loss of AcrB function (6).

Other transcriptomic studies identified genes in group one being as being differentially regulated in the *S*. Typhimurium wild type and strains lacking *acrB* or *acrB* function (*acrB* D408A mutant). Comparison in exponential phase in MOPS minimal medium (5, 6) revealed that SPI-1 genes were downregulated in an *acrB* knockout mutant and in the *acrB* D408A mutant. In stationary phase (7), rRNAs, tRNAs, SPI-11 and some SPI-1 genes were downregulated (although the SPI-1 *sipABCD* TTSS genes, not members of groups one-three in the present study, were upregulated in the mutant). However, our data suggests that this is a response to fundamental changes in physiology brought about by deletion of *acrB*, and not necessarily direct regulation of pathogenicity islands.

### Group two – early activation in the ΔacrB strain

The 46 genes in group two (Supplemental table S2) are more upregulated between 1 h and 3 h in the *acrB* strain than in the same time period in the wild type, so are classified as “early activation”. On initial inspection, this group contains many genes known to be anaerobically activated. For each gene, expression data in the gene expression database SalComMac (22, 23) was consulted to identify which genes are anaerobically activated; in this database, “anaerobic shock” is classified as aerobic growth followed by 30 minutes in a sealed static tube (22). It should be noted that the SalComMac data is for *S*. Typhimurium strain 4/74; SL1344 is a *hisG* auxotroph of 4/74 (24), although we believe that data should be broadly comparable for the two strains.

Many of the anaerobically-activated genes are activated by the oxygen-sensing global regulator FNR in *Salmonella* (25) and / or *E. coli* (26). These include genes encoding the periplasmic nitrate and nitrite reductases responsible for anaerobic respiration at low NO_2_^-^ and NO_3_^-^ concentrations (27, 28), the anaerobic serine/threonine degradation enzyme TdcB that is suggested to be critical for energy metabolism during changes in oxygen tension (29), and the two cytochrome *c* maturation (Ccm) operons, one of which is downstream of the *nap* operon (30). The cytochrome *c* peroxidase *yhjA*/*ccp* is also in this group, which has been shown to be FNR-activated (25) and enables respiratory growth on peroxide (31). The *ccp* gene is upstream of the second *ccm* cluster and may form a single operon. RS22105, annotated as a molybdopterin-dependent oxidoreductase and proposed to be a DMSO reductase distinct from DmsABC, has been shown to be FNR activated in *Salmonella* (25), as has *yhbU/ubiU* which encodes part of the oxygen-independent ubiquinone synthesis pathway (32).

A high affinity nickel uptake system (*yntABCDE/nikABCDE*) is in the grouping (33); these genes are FNR activated in *E. coli* although were not identified in the *Salmonella* FNR study (25). In *E. coli, nikABCDE* expression is linked to that of the Ni^2+^ cofactor-containing hydrogenases required for respiration of fumarate and formate (34). RS15890 and RS15900 (*ecfT*) are two members of a putative 5-gene operon encoding an ECF transporter suggested to import Co^2+^ (35) that is conserved in pathogenic *E. coli* strains. SalComMac data suggests that this operon is anaerobically activated.

The remaining anaerobically-activated genes in this group are of unknown function or are poorly characterised: *yecH* has been shown to be downregulated by adrenaline (36); *ompX* is in SPI-5 (37) and has been found to be overexpressed in a *dksA* mutant (38); two LysR family transcriptional regulators (RS11710 and RS19740); a putative transketolase RS12010; and hypothetical protein RS12010.

It should be noted that for many of these anaerobic genes, expression increases from 1 h to 3 h and then decreases from 3 h and to 5 h. The magnitude of the increase in expression is larger in the *acrB* strain. Taken together, genes in this group enable anaerobic energy generation through respiratory and substrate-level routes.

As well as the anaerobically-activated genes in this group, there are also several genes that are anaerobically repressed in SalComMac, including the tyrosine importer *tyrP*, the flagellar genes *fliR, fliF* and *fliH* and the proposed virulence factor *srfA* which was also identified as a class 2 flagellar gene (39) whose expression is activated by the flagellar master regulator FlhDC. As previously noted, loss of AcrB function reduces *Salmonella* motility (5). A link between motility and SPI-1 has also been reported (40).

In previous transcriptomic studies, Wang-Kan et al (6) identified anaerobically-activated genes including nitrate and nitrite reductase and flagellar genes being upregulated in the *acrB* D408A mutant in exponential phase. In stationary phase, some anaerobically-relevant and flagellar genes were also upregulated (7).

### Group three – late activation in the ΔacrB strain

Group three comprises 53 genes that are more activated in the wild type than in Δ*acrB* at three hours, so the opposite pattern to genes in groups one and two (Supplemental Table S3). The most striking group of genes in this group are the *met* genes involved in methionine biosynthesis from aspartate, methionine uptake, and S-adenosyl methionine (SAM) biosynthesis; expression of these genes is seen to “spike” at 3 h in the wt, with upregulation at 3 h vs 1 h and downregulation at 5 h vs 3 h. In the Δ*acrB* strain these genes are less upregulated at 3 h and many are further upregulated at 5 h, indicating much later induction. In *E. coli* these genes are all repressed by the regulator MetJ which binds to methionine and SAM (41). Our data suggest that these routes to SAM are highly upregulated on entry to stationary phase. SAM is an essential metabolite and a major methyl donor for protein and DNA methylation and polyamine synthesis; in the transition to stationary phase, SAM is also a methyl donor for cyclopropane fatty acid synthesis (42). In *E. coli*, the SAM synthase *metK* is essential (43) and depletion of SAM using a *metK84* mutant leads to a cell division defect (44). In *S*. Typhimurium, methionine biosynthesis or import are required for infection in mice and methionine was implicated in aspects of peptidoglycan synthesis (45).

These data suggest that the methionine and / or SAM pool in the wild type is depleted at 3 h growth, and these pathways are upregulated to replenish the pool. The transcriptomics data in SalComMac shows that most of the *met* genes are not upregulated in stationary phase, although that dataset was generated for cultures grown in rich Lennox broth (22). We suggest that in Lennox broth, methionine is available (around 70 mg/L based on typical composition values (46)) so can be readily imported and converted to SAM, whereas in MOPS minimal medium used here, methionine needs to be synthesised from aspartate, hence activation of the *met* genes. This suggests that methionine could be a growth limiting nutrient in these conditions.

A cluster of genes (*ybdL, ybdH, ybdD, ybdM*) are also regulated in a similar manner to the *met* genes. *E. coli* YbdL has methionine aminotransferase activity (47) and is believed to be involved in methionine recycling following polyamine synthesis (48). YbdH is annotated as a putative glycerol dehydrogenase and the homologue in *E. coli* has been identified as a hydroxycarboxylate dehydrogenase (49). The *E. coli ybdD* homologue has been identified (with CstA) as part of a pyruvate import system (50) and *ybdM* has been identified as a transcriptional regulator related to SpoOJ (51). A predicted MetJ binding site in this cluster could explain its regulation (41).

The PstSCAB high-affinity phosphate importer is poorly understood in *Salmonella* but is known to be induced in *E. coli* in response to phosphate limitation, and is activated by RpoS (52). ProP is responsible for the uptake of osmoprotectants proline and glycine betaine; expression in *E. coli* is induced by RpoS and Fis on entry to stationary phase (53). Expression of *mysB, yahO*, and *yceK* are also RpoS-induced (54). Expression of *katG* in *E. coli* was also shown to increase on entry into stationary phase, although in an RpoS-independent manner (55).

Two genes with key roles in regulating entry into stationary phase entry in this group: *rsd* encodes an anti-sigma factor that deactivates the exponential phase housekeeping sigma factor σ^70^ RpoD (56); and *rmf* induces ribosome dimerization and hibernation (57). Expression of *rmf* is highly upregulated in stationary phase in the SalComMac dataset (22). Both of these genes are upregulated far later in the *acrB* mutant than in the wild type in our dataset; together with the other stationary-phase induced genes in group three, this supports the later onset of stationary phase in Δ*acrB* than the wild type.

Expression of *aceB* is regulated by multiple transcription factors in *E. coli*, including ArcA (58). There are also a number of genes in SPI-2, involved in intracellular survival, and chemotaxis-related genes in group three.

Lower expression of SPI-2 genes were observed in other transcriptomics studies in the *acrB* D408A mutant in exponential (6) and stationary phase (7), although the *met* genes and many others in group three were not reflected in the datasets of previous studies. The reason for this is likely because most genes in this group were transiently expressed in the wild type; we observed a “spike” of expression, and later expression in the Δ*acrB* mutant. This reveals a key advantage of our approach; such a “spike” could be missed if only a single timepoint was used to measure transcriptomes. Taking samples through growth allows the expression of each gene to be tracked and the profiles of gene expression over time to be compared.

### A model linking efflux and stationary phase via membrane potential

Previous studies have proposed that changes in gene expression and physiology in efflux knockouts have been caused by the lack of efflux of a signalling molecule (for example an autoinducer), giving rise to changes in intracellular and/or extracellular concentrations of that molecule and a resultant change in gene regulation and physiology. However, we realised that knockout of *acrB* could trigger an alternative signalling cascade. As AcrB requires proton motive force (PMF) for function, deactivation of AcrB would decrease of influx of protons across the inner membrane, resulting in membrane hyperpolarisation and thus alter the redox state of the cell. This has been built into a model (Figure 3) whereby the membrane potential and redox state of the bacterium plays a key role in the transition from exponential to stationary phase, mediated by the two-component ArcBA system (59). Deletion of AcrB results in perturbation of the membrane potential and thereby the redox balance of the cell and thus disrupts this regulation, resulting in a) late entry into stationary phase and b) upregulation of anaerobic energy metabolism pathways.

**Figure 3.**
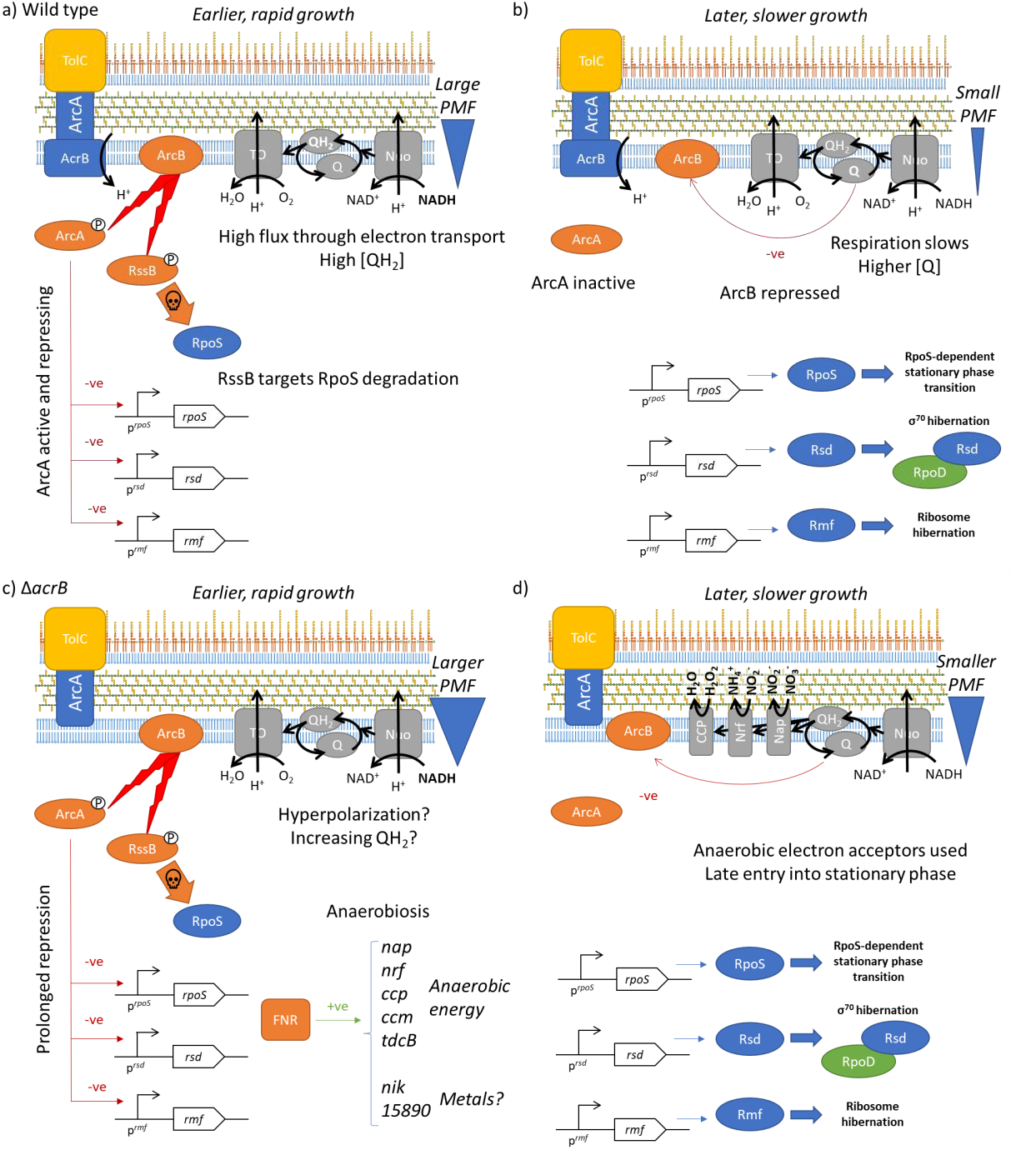
Model of the mechanism of ArcBA-mediated redox control of stationary phase entry and the impact of an AcrB mutant. In the wild type strain during rapid growth (a), high flux through the electron transport chain results in a reduced quinol pool (QH_2_) which permits ArcBA to be active, repressing expression of *rpoS, rsd* and *rmf*. RssB also targets RpoS protein for degradation. As growth slows (b), the quinol pool becomes more oxidised (Q), deactivating ArcB, permitting activation of *rpoS, rsd* and *rmf*, triggering entry into stationary phase. In a *ΔacrB* mutant (c), proton motive force is higher for longer during growth, repressing *rpoS, rsd* and *rmf* and delaying entry into stationary phase. As a result, exponential phase physiology is extended (d), necessitating upregulation of genes involved in anaerobic energy metabolism (Nap, Nrf and CCP shown), likely via the global regulator FNR. TO: Terminal oxidases.

The global regulator ArcBA regulates gene expression on the shift from aerobic to anaerobic metabolism (59) and is also thought to play a role in response to other stresses such as reactive oxygen species (ROS) and during infection in *Salmonella* (60, 61). Knockouts of *arcA* and *arcB* in *S*. Typhimurium reduce intracellular survival in epithelial cells, macrophages and neutrophils, and are attenuated in a mouse model (60). The ArcBA regulon has been characterised in *E. coli* (reviewed by (59)) and *Salmonella* both aerobically and anaerobically (61, 62). The sensor kinase ArcB autophosphorylates itself and then phosphorylates the response regulator ArcA; phospho-ArcA binds promoter DNA and regulates gene expression (Figure 3a). Oxidised quinones repress the kinase activity of ArcB (63) thus ArcBA is a direct sensor of the quinol pool redox state. Highly oxidised quinone (Q) leads to low ArcA activity whereas highly reduced quinol (QH_2_) leads to high ArcA activity.

ArcA has previously been shown to repress expression of *rpoS, rsd*, and *rmf*, three genes that mediate important aspects of the switch from exponential phase to stationary phase (14, 64). RpoS is the sigma factor that is dominant in stationary phase (14); *rpoS* transcription is repressed by ArcA in *E. coli* (65) and *Salmonella* (66). In addition, activated ArcB phosphorylates RssB, which targets RpoS protein for degradation by the ClpXP protease (65). ArcAB-RssB therefore acts as a key gatekeeper to entry into stationary phase. Rsd is an anti-sigma factor that deactivates the exponential phase housekeeping sigma factor σ^70^ / RpoD (56). Expression of *rsd* is regulated by ArcA in both *E. coli* (67) and *Salmonella* (61). Rmf induces ribosome dimerization and hibernation in stationary phase (57); its expression is ArcA-regulated in *Salmonella* (61) and *E. coli* (64).

In the wild type in rapid growth, relatively high flux through electron transport to oxygen results in a highly reduced quninol pool (QH_2_). This would enable ArcBA to be active, repressing *rpoS, rsd* and *rmf*. During entry into stationary phase, the quinol pool becomes more oxidised, de-activating ArcBA and thus permitting *rpoS, rsd* and *rmf* expression and activity. In the transition of the *ΔacrB* strain to stationary phase, the inner membrane would become hyperpolarised; this would alter the redox state of the quinol pool, meaning that ArcBA remains active and RpoS, Rsd and Rmf activities are repressed for longer during growth. This would delay onset of stationary phase in the Δ*acrB* mutant. The genes *rsd* and *rmf* both fall into group three; Table 1 summarises their expression data. Expression of *rpoS* is insignificant (*p*_*adj*_ > 0.05) in the wild type between 1 h and 3 h, although downregulated between 1 h and 3 h in the Δ*acrB* mutant.

**Table 1.** Gene expression data for the key stationary phase regulators *rsd, rmf*, and *rpoS*. Log_2_ fold change values for each gene: 1 h vs 3 h in the wild type; 3 h vs 5 h in the wild type; 1 h vs 3 h in Δ*acrB*; 3 h vs 5 h in Δ*acrB*. ΔLog_2_ values for the 1 h vs 3 h and 3 h vs 5 h comparisons. Δlog_2_ values are calculated by Log_2_ fold change for Δ*acrB* minus Log_2_ fold change for the wild type. ID: locus name. Fold changes that are insignificant (*p*_*adj*_ > 0.05) are marked with a dagger †. Other fold changes are colour coded as in Supplemental tables.

| Log<sub>2</sub> fold change | | | | Δ*acrB* – wt<br>Δlog<sub>2</sub> | | ID | Gene | STM |
| --- | --- | --- | --- | --- | --- | --- | --- | --- |
| wt 1vs3 | wt 3vs5 | Δ*acrB* 1vs3 | Δ*acrB* 3vs5 | 1vs3 | 3vs5 | | | |
| 0.98 | 0.35<sup>†</sup> | -0.69 | 1.59 | -1.67 | 1.24 | SL1344_RS21380 | *rsd* | STM4165 |
| 0.36<sup>†</sup> | 0.73<sup>†</sup> | -1.17 | 2.19 | -1.53 | 1.46 | SL1344_RS15130 | *rpoS* | STM2924 |
| 2.30 | 3.74 | 1.05 | 4.61 | -1.25 | 0.87 | SL1344_RS05235 | *rmf* | STM1066 |

In the absence of a transition into stationary phase, exponential phase physiology would be prolonged in the Δ*acrB* mutant with σ^70^ remaining the dominant sigma factor. Oxygen limitation caused by increased biomass concentration would likely activate FNR, which would upregulate the anaerobic energy-generating pathways identified in group two above. Nrf, Nap and CCP do not pump protons across the inner membrane (68), therefore they would not contribute to the hyperpolarised state of the bacterium. ArcA does not appear to regulate the anaerobically-activated genes identified here in Supplemental Table S2 except for the periplasmic nitrite reductase *nrf* (62).

### Experimental determination of membrane potential

To test whether the inner membrane of the Δ*acrB* strain is hyperpolarised, we used the fluorescent membrane potential dye DiSC_3_(5) (3,3′-dipropylthiadicarbocyanine iodide) (69). The wild type, Δ*acrB* and *acrB* D408A strains were grown in MOPS minimal medium and the membrane potential measured using flow cytometry after 1, 3 and 5 hours (Figure 4). As a control, the protonophore carbonyl cyanide m-chlorophenylhydrazone (CCCP) which collapses the PMF was used prior to measurement. The wild type strain showed highest membrane potential after 1 hours growth, and declining membrane potential at 3 and 5 hours. Both the Δ*acrB* strain and a strain expressing the *acrB* D408A derivative, which blocks proton translocation and thus efflux (70, 71), had higher membrane potential than the wild type at all time points. These data support the hypothesis that the Δ*acrB* mutant is hyperpolarised. Treatment with CCCP collapsed the membrane potential of all strains at all time points (*P* ≤ 0.01).

**Figure 4.**
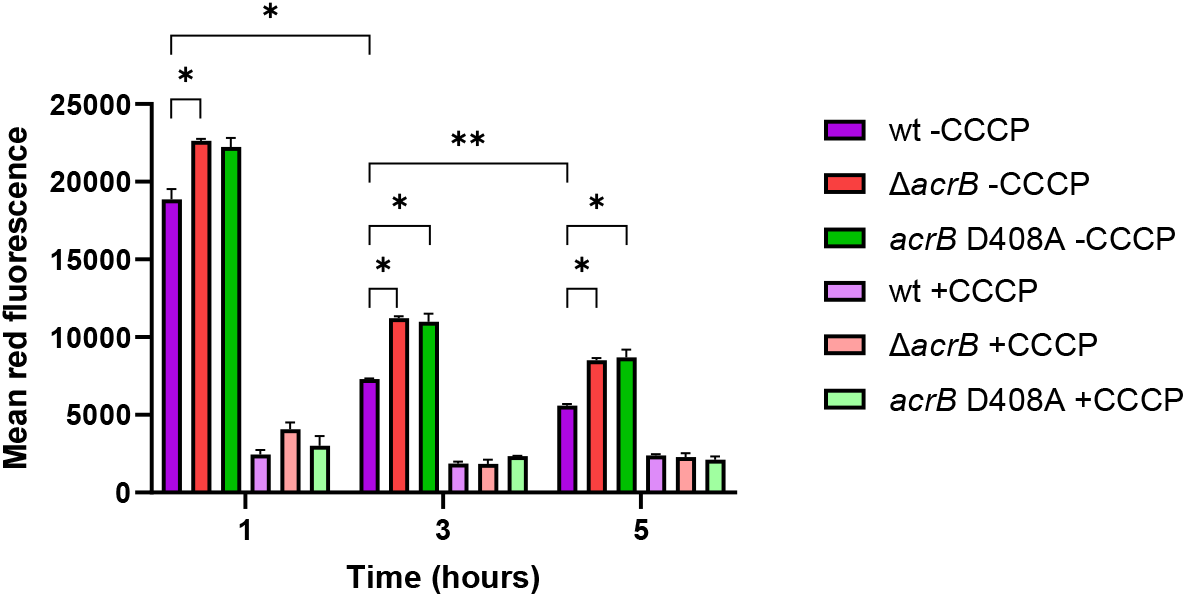
Determination of membrane potential. SL1344, Δ*acrB* and *acrB* D408A cells were grown in MOPS minimal medium and samples taken at 1, 3 and 5 hours growth, whereupon membrane potential was determined using flow cytometry and DiSC_3_(5). The red fluorescence, corresponding to membrane potential, is shown for each strain. The protonophore CCCP was used as a control to collapse membrane potential. Error bars show standard deviation of duplicate cultures for a representative experiment. Significance was tested for comparisons of wild type and Δ*acrB* / *acrB* D408A strains for each timepoint and across timepoints using a paired t-test; * *P* ≤ 0.05; ** *P* ≤ 0.01. Addition of CCCP to each sample resulted in a significant (*P* ≤ 0.01) decrease in membrane potential.

There is only one other study where the influence of efflux pumps on membrane potential has been measured; Le *et al*. showed that PMF was similar in wt and *tolC* strains of *E. coli* by measuring flagellar rotor speed (72). These experiments were done on cells grown in rich medium to an optical density of 2, so are not directly comparable with our data.

## Discussion

Our transcriptomics data suggests that the Δ*acrB* strain enters stationary phase later than the wild type, and as a result displays prolonged exponential-phase anaerobic metabolism. Comparison with other transcriptomic experiments on efflux pump mutants reveals some similarities in expression patterns. Wang-Kan *et al*. compared expression in SL1344 and an efflux-defective *acrB* D408A mutant (6). The two strains were grown in MOPS minimal medium to an OD_600_ of 0.6. The *acrB* D408A mutant displayed higher expression of flagella and anaerobic metabolism genes than the wild type. A later transcriptomic study (7) grew the same strains in LB broth to stationary phase and found changes in genes responsible for energy metabolism. A major advantage of measuring gene expression across growth (here at 1, 3 and 5 hours) is that trends in gene expression (and thereby physiology) over time can be identified, which would otherwise be difficult to detect or interpret when taking samples at only a single timepoint.

### Supporting evidence for the redox-ArcBA model from the literature

There is data in the literature to support the model of efflux pump function influencing membrane potential and regulation of stationary phase (Fig. 3). The links between efflux pumps and membrane potential have previously been observed in studies on efflux pump inhibitors. The antihistamine promethazine (a phenothiazine) is an efflux pump inhibitor (73) and has been shown to increase membrane potential in *Pseudomonas aeruginosa* (74).

Arce-Rodríguez *et al*. studied the impact of the NADH:NAD^+^ ratio on antibiotic resistance in *Pseudomonas aeruginosa* PA14 (75). They artificially modulated the NADH:NAD^+^ ratio using a recombinant NADH oxidase (which increases [NAD^+^]) or a formate dehydrogenase (which increases [NADH]). They showed that a high NADH:NAD^+^ ratio increased membrane potential whereas low NADH:NAD^+^ ratios decreased membrane potential (measured using DiOC_2_(3)). High NADH:NAD^+^ ratios also increased cellular ROS levels, and efflux of aminoglycosides and fluoroquinolones via the RND efflux pumps MexAB and MexXY leading to increased resistance. However, a high NADH:NAD^+^ ratio also increased ROS-mediated killing by high concentrations of antibiotics, and it was concluded that control of NADH:NAD^+^ ratio and other aspects of redox balance was important for response to antibiotics.

Yang *et al*. found that an Δ*acrA* mutant in *E. coli* BW25113 had lower *rpoS::lacZ* expression than the wild type across growth, while overproduction of AcrA resulted in increased *rpoS::lacZ* expression (76). Yang *et al*. proposed that efflux pumps export quorum sensing molecules which gives rise to these changes in physiology, but we present an alternative model whereby membrane potential and redox state is the signal. Indeed, subsequent studies have sought to identify effluxed quorum sensing molecules without success (7). A previous study found that *rpoS* transcription in *S*. Typhimurium F98 is linked to redox and influenced by oxygen, ArcA, and NADH:NAD^+^ ratio (66). Expression of *arcA* was also found to be negatively regulated by RpoS. The authors also suggested that quorum sensing could play a role in this regulation. ArcA has also recently been found to be important in *E. coli* for development of antibiotic resistance (77) and metabolic adaptation to tetracycline resistance (78).

### Links to biofilms?

It has previously been reported that knockout or deactivation of *acrB* and other genes encoding efflux pump components reduced biofilm formation in *Salmonella* (79), although the exact regulatory mechanism is currently unclear (80, 81). Indeed, efflux pumps have been implicated in biofilm formation in many bacterial species (reviewed by (82)). A recent study identified TolC as being beneficial for biofilm formation in both *E. coli* and *S*. Typhimurium (81). This study also identified *nuo* genes encoding the major aerobic NADH reductase as being important in *Salmonella* biofilm formation, and highlighted the fact that biofilm development is a process requiring different genes at different times (from initial attachment to maturation). Efflux inhibitors such as phenothiazines have also been shown to reduce biofilm formation (73, 74). ArcBA and RpoS have previously been implicated in regulation of biofilm formation (83) so our model could provide a link between efflux pumps, redox state, and biofilm formation. Indeed, recent studies have shown that *Salmonella* biofilm formation can be modulated by redox-active polymers (84).

## Conclusions

In summary, we have demonstrated that deactivation of AcrB leads to hyperpolarisation of the inner membrane and we propose that this is responsible for wide-ranging changes in bacterial gene regulation and physiology. Our model linking efflux and redox provides an explanation for some phenotypes observed in efflux pump knockouts and inhibition experiments, and places efflux pumps as a central part of bacterial physiology, distinct from (but linked to) their role as transporters of noxious substances from the cell.

Intriguingly, our model could explain three reasons why efflux pumps are important for the response of *Salmonella* to antimicrobials. First, their well-established and characterised function in removing noxious substances from the cell. Second, as loss of AcrB function increases membrane potential, antimicrobials postulated to function via the generation of reactive oxygen species by the respiratory chain following treatment (the ROS-lethality hypothesis) (85) could be more effective. And third, as the Δ*acrB* strain appears to enter stationary phase later in growth than the wild type, inhibition of efflux is likely to delay the remodelling of the cell envelope that occurs on transition to stationary phase, after which antimicrobial permeability is far reduced (9). Likewise, the attenuation of virulence observed when AcrB is deactivated (6) could be due to deficiencies in efflux of antimicrobial host molecules, and / or changes in physiology brought about by hyperpolarisation such as alterations in SPI expression, motility, and biofilm formation. Future work will explore the relative contributions of efflux and redox state in these situations.

## Materials and Methods

*Salmonella enterica* serovar Typhimurium SL1344 (86) and its cognate Δ*acrB* (87) and *acrB* D408A mutant (6) were used. Transcriptomic analysis on SL1344 and SL1344 Δ*acrB* was performed as described by (9). RNA-Seq data have been deposited with Array Express (accession no. E-MTAB-9679).

Total numbers of differentially expressed genes (DEGs) were determined using the R package DESeq2 (88) by comparing the three timepoints (1h, 3h, and 5h) for both the wild type and the Δ*acrB* strain. Adjusted p-value < 0.05 and log_2_FC ≥ |1.5| were both used as the thresholds for determining the DEGs for the Venn diagrams (Figure 1), which were drawn using the R package ggvenn (89).

Genes were classified in groups 1, 2 and 3 by the following criteria. The log_2_ fold change for each gene was calculated for 1 h versus 3 h in both wt and Δ*acrB* strains. Only genes where log_2_FC data were significant (p_*adj*_ < 0.05) for the 1 h vs 3 h comparison for both wt and Δ*acrB* strains were considered for this analysis. Δlog_2_FC was calculated for each gene by subtracting the log_2_FC in the wt from the log_2_FC in the Δ*acrB* strain. Groups were classified: group 1, wt log_2_FC for 1 h vs 3 h was ≤ -1 and Δlog_2_FC was ≥ 1; Group 2, wt log_2_FC for 1 h vs 3 h was > -1 and Δlog_2_FC was ≥ 1; Group 3, Δlog_2_FC was ≤ 1.

### Membrane potential measurement by flow cytometry

Strains SL1344, Δ*acrB* and *arcB* D408A were grown in 10 mL of MOPS minimal medium (9) at 37 °C. After 1, 3 and 5 hours, cells were harvested by centrifugation (4720 x g for 5 min) and resuspended in filter-sterilised HEPES-buffered saline (HBS, Alfa Aesar). Diluted cell suspensions were added to tubes containing HBS. 15 μM CCCP was added to control samples and incubated for 15 min with shaking in the dark. To all samples was added 1 μM DiSC_3_(5), 1% DMSO, 0.2% Glucose, and 1 mM CaCl_2_ before being shaken for a further 15 min in the dark (69). Samples were then analysed using a C6 Plus flow cytometer (BD Biosciences) and CFlow plus software. Data rate was 1000 to 4000 events/sec and a FSC-H threshold of 8000 was used to discriminate noise from bacteria. Bacteria were gated using forward scatter and side scatter measurements from blue laser illumination (488 nm), and red fluorescence was measured using red laser illumination (640 nm) and collected using a 670 LP filter.

## Author contributions

J.M.A.B., T.W.O., and E.E.W. designed the assays. E.E.W. performed experiments to obtain samples for RNA-seq. A.O. and S.J.E. performed experiments to measure membrane potential and analysed the resulting data. RNA-seq data were analyzed by E.E.W., O.O., and T.W.O. The manuscript was written by J.M.A.B. and T.W.O. with input from all authors.

