## Supplemental information for "Efflux pumps mediate changes to fundamental bacterial physiology via membrane potential"

**Contents:**

Supplemental Table S1 2

Supplemental Table S2 9

Supplemental Table S3 12

Supplemental References 15

### Supplemental Table S1. Genes in group one - Genes that are less downregulated at 3hr in Δ*acrB* than wild type – "late repression" pattern

| Log$_2$ fold change | | | | Δ*acrB*-wt Δlog$_2$ | | | | | | | | |
|---|---|---|---|---|---|---|---|---|---|---|---|---|
| wt 1vs3 | wt 3vs5 | acrB 1vs3 | acrB 3vs5 | 1vs3 | 3vs5 | ID | Gene name | Operon structure | STM | Product | Comments | Regulation in *E. coli* |
| -4.29 | 0.00 † | -0.53 | -3.94 | 3.77 | -3.94 | RS12415 | tRNA0043 | tRNA0040-43 | | tRNA-Lys (ttt) | tRNA & related | |
| -3.53 | 0.82 | 0.22 | -2.95 | 3.75 | -3.77 | RS13050 | *suhB* | *suhB* | STM2546 | Nus factor; inositol-1-monophosphatase | Translation | |
| -4.08 | -0.65 † | -0.36 | -4.08 † | 3.73 | -3.43 | RS19775 | *yidD* | *rpmHA-yidDC* | STM3841 | membrane protein insertion efficiency factor | Envelope – IM protein insertion | |
| -3.96 | 0.22 † | -0.36 | -3.26 | 3.60 | -3.47 | RS03415 | tRNA0010 | tRNA0013-07 | | tRNA-Gln (ttg) | tRNA & related | |
| -4.31 | 0.04 † | -0.93 | -3.35 | 3.37 | -3.39 | RS03420 | tRNA0011 | tRNA0013-07 | | tRNA-Gln (ttg) | tRNA & related | |
| -3.86 | -0.51 † | -0.54 | -3.07 | 3.32 | -2.56 | RS10300 | tRNA0033 | tRNA0033 | | tRNA-Asn (gtt) | tRNA & related | |
| -3.75 | 0.27 † | -0.55 | -2.96 | 3.20 | -3.23 | RS03430 | tRNA0013 | tRNA0013-07 | | tRNA-Met (cat) | tRNA & related | |
| -3.58 | 0.20 † | -0.43 | -3.28 | 3.15 | -3.47 | RS20225 | tRNA0067 | tRNA0067 | | tRNA-His (gtg) | tRNA & related | |
| -3.44 | -0.13 † | -0.43 | -3.25 | 3.01 | -3.11 | RS12410 | tRNA0042 | tRNA0040-43 | | tRNA-Val (tac) | tRNA & related | |
| -2.45 | 0.87 | 0.40 | -2.22 | 2.85 | -3.09 | RS22345 | tRNA0080 | tRNA0080-82 | | tRNA-Gly (gcc) | tRNA & related | |
| -2.18 | 0.51 † | 0.67 | -2.19 | 2.84 | -2.70 | RS04610 | tRNA0019 | tRNA0019 | | tRNA-Ser (gga) | tRNA & related | |
| -3.29 | -0.17 † | -0.47 | -3.39 | 2.82 | -3.22 | RS14630 | tRNA0050 | tRNA0050-46 | | tRNA-Ser (gct) | tRNA & related | |
| -2.37 | 0.76 | 0.41 | -1.90 | 2.78 | -2.66 | RS22355 | tRNA0082 | tRNA0080-82 | | tRNA-Gly (gcc) | tRNA & related | |
| -3.33 | 0.01 † | -0.55 | -2.72 † | 2.78 | -2.73 | RS16590 | *rpsU* | *rpsU* | STM3029 | 30S ribosomal protein S21 | Ribosome | ppGpp-DksA, LexA repressed |
| -3.37 | 0.25 † | -0.72 | -2.74 | 2.64 | -3.00 | RS01270 | tRNA0002 | tRNA0002 | | tRNA-Ala (tgc) | tRNA & related | |
| -3.08 | 0.34 † | -0.44 | -2.78 | 2.64 | -3.12 | RS20220 | tRNA0066 | tRNA0066-69 | | tRNA-Arg (ccg) | tRNA & related | |
| -3.13 | 0.83 | -0.50 | -1.71 | 2.63 | -2.54 | RS15255 | *queD* | *queD* | STM2949 | 6-carboxytetrahydropterin synthase | tRNA & related | Nac repressed |
| -3.29 | 0.16 † | -0.68 | -2.17 | 2.61 | -2.33 | RS20500 | tRNA0070 | tRNA0070 | | tRNA-Ile (gat) | tRNA & related | |
| -3.19 | -0.48 † | -0.59 | -2.74 | 2.60 | -2.26 | RS01265 | tRNA0001 | tRNA0001 | | tRNA-Ile (gat) | tRNA & related | |
| -3.24 | 0.20 † | -0.66 | -2.13 | 2.58 | -2.32 | RS06145 | *cspF* | *cspF* | STM1243 | Cold shock-like protein | SPI-11 | |
| -1.88 | 1.30 | 0.67 | -1.72 | 2.55 | -3.02 | RS13200 | *yfhL* | *yfhL* | STM2576 | Putative ferrodoxin | Unknown | Nac repressed |
| -3.19 | 0.08 † | -0.68 | -2.22 | 2.51 | -2.30 | RS21225 | tRNA0067 | tRNA0066-69 | | tRNA-Ile (gtg) | tRNA & related | |
| -3.33 | 0.46 † | -0.84 | -2.39 | 2.49 | -2.85 | RS21230 | tRNA0068 | tRNA0066-69 | | tRNA-Ala (cag) | tRNA & related | |
| -2.85 | 1.62 | -0.40 | -1.17 | 2.45 | -2.78 | RS04505 | *artP* | *artPJQM* | STM0891 | arginine ABC transporter ATP-binding protein ArtP | AA metabolism - Arginine uptake | ArgR repressed, Lrp regulated |

| | | | | | | | | | | | | |
|---|---|---|---|---|---|---|---|---|---|---|---|---|
| -3.01 | -0.49 † | -0.57 | -2.70 | 2.44 | -2.21 | RS04630 | *infA* | *infA* | STM0953 | translation initiation factor IF-1 | Translation | ppGpp repressed |
| -3.13 | -0.69 | -0.70 | -2.97 | 2.43 | -2.28 | RS05850 | *rluC* | *rluC* | STM1187 | 23S rRNA pseudouridine(955/2504/2580) synthase RluC | Ribosome | ppGpp repressed |
| -1.60 | 0.00 † | 0.81 | -2.98 | 2.41 | -2.98 | RS16610 | tRNA0055 | tRNA0055 | | tRNA-Ile (cat) | tRNA & related | |
| -3.24 | 0.16 † | -0.86 | -2.37 | 2.39 | -2.53 | RS20505 | tRNA0071 | tRNA0071 | | tRNA-Ala (tgc) | tRNA & related | |
| -3.21 | 0.75 | -0.83 | -1.70 | 2.38 | -2.45 | RS07340 | *ydgI* | *ydgI* | STM1477 | Putative arginine:ornithine antiporter | AA metabolism - Arginine import | ArgR repressed |
| -2.67 | -0.09 † | -0.39 | -2.08 | 2.28 | -1.99 | RS08775 | *lolB* | *lolB-ispE* | STM1778 | Lipoprotein localization protein LolB | Envelope – OM lipoprotein trafficking | |
| -2.83 | -0.43 † | -0.58 | -2.72 | 2.25 | -2.28 | RS09715 | tRNA0028 | tRNA0030-28 | | tRNA-Leu (taa) | tRNA & related | |
| -3.04 | -0.33 † | -0.82 | -2.44 | 2.22 | -2.10 | RS04235 | *ldtB* | *ldtB* | STM0837 | L'D-transpeptidase | Envelope – Peptidoglycan remodelling | Rob activated, CRP regulated |
| -2.49 | -2.18 | -0.29 | -4.11 † | 2.20 | -1.93 | RS27065 | *yoeI* | *yoeI-plaP* | | Membrane protein | Unknown | |
| -2.75 | -0.39 † | -0.60 | -2.43 | 2.14 | -2.04 | RS23375 | *rsmC* | *rsmC* | STM4556 | 16S rRNA (guanine(1207)-N(2))-methyltransferase RsmC | Ribosome | |
| -2.83 | 0.62 † | -0.70 | -1.80 | 2.13 | -2.42 | RS06595 | *ydiY* | *ydiY* | STM1327 | YdiY family protein, putative OMP | Unknown | Nac repressed |
| -2.75 | -0.06 † | -0.62 | -2.35 | 2.13 | -2.29 | RS09720 | tRNA0029 | tRNA0030-28 | | tRNA-Cys (gca) | tRNA & related | |
| -2.72 | 0.11 † | -0.59 | -2.07 | 2.13 | -2.18 | RS09725 | tRNA0030 | tRNA0030-28 | | tRNA-Gly (gcc) | tRNA & related | |
| -2.61 | -0.55 † | -0.55 | -2.01 | 2.06 | -1.46 | RS24830 | RtT RNA | | | RtT sRNA | Regulation | |
| -2.52 | 1.59 | -0.47 | -0.96 | 2.05 | -2.54 | RS02070 | *queA* | *queA* | STM0404 | tRNA preQ1 (34) S-adenosylmethionine ribosyltransferase-isomerase QueA | tRNA & related | |
| -1.11 | 0.31 † | 0.88 | -1.64 | 1.99 | -1.95 | RS10975 | *trhP (yegQ)* | *trhP* | STM2136 | tRNA wobble base hydroxylation protein TrhP | tRNA & related | |
| -2.27 | -1.70 | -0.29 | -3.65 | 1.98 | -1.95 | RS19205 | *rpmB* | *rpmB* | STM3728 | 50S ribosomal protein L28 | Ribosome | ppGpp / DksA repressed |
| -2.34 | -1.78 | -0.39 | -3.73 † | 1.96 | -1.95 | RS19765 | *rpmH* | *rpmHA-yidDC* | STM3839 | 50S ribosomal protein L34 | Ribosome | ppGpp repressed |
| -2.82 | -1.06 | -0.89 | -3.01 | 1.94 | -1.95 | RS04740 | *ycaO* | *ycaO* | STM0975 | 30S ribosomal protein S12 methylthiotransferase accessory protein YcaO | Ribosome | |
| -2.81 | -0.73 | -0.88 | -2.25 | 1.93 | -1.52 | RS11365 | *yeiU (lpxT)* | *yeiRU* | STM2213 | Phosphatase PAP2 family protein | Envelope - Lipid A modification | |
| -2.47 | -0.74 † | -0.55 | -2.95 | 1.92 | -2.21 | RS17070 | *rplU* | *rplU-rpmA* | STM3304 | 50S ribosomal protein L21 | Ribosome | ppGpp / DksA, Nac repressed; MlrA activated |
| -1.22 | -0.49 † | 0.69 | -1.68 | 1.91 | -1.19 | RS23015 | tRNA0083 | tRNA0083 | | tRNA-Leu (caa) | tRNA & related | |
| -2.32 | 1.21 † | -0.48 | -0.43 † | 1.84 | -1.64 | RS03590 | *dtpD* | *dtpD* | | Putative MFS transporter, homology to *E. coli* dipeptide importer *dtpD* | Transport | |
| -2.12 | 0.03 † | -0.27 | -1.79 | 1.84 | -1.82 | RS04310 | *rimO* | *rimO* | STM0852 | 30S ribosomal protein S12 methylthiotransferase accessory protein RimO | Ribosome | |
| -2.41 | -1.63 | -0.59 | -3.52 † | 1.82 | -1.89 | RS00220 | *rpsT* | *rpsT* | STM0043 | 30S ribosomal protein S20 | Ribosome | ppGpp / DksA repressed |
| -2.37 | -0.70 | -0.60 | -2.19 | 1.78 | -1.49 | RS17005 | *secG* | *secG-* tRNA0057 | STM3293 | Preprotein translocase subunit | Envelope - Protein translocation | Nac repressed |
| -2.97 | -0.41 † | -1.20 | -2.11 | 1.77 | -1.70 | RS19950 | *mioC* | *mioC* | STM3875 | FMN-binding protein | Cell division | |
| -2.46 | -0.20 † | -0.70 | -2.15 | 1.76 | -1.94 | RS04765 | *cmk* | *cmk* | STM0980 | Cytidylate kinase | Nucleotide metabolism - Pyrimidine salvage | |
| -2.46 | -1.17 | -0.70 | -2.10 | 1.76 | -0.93 | RS22135 | RS22135 | RS22150-22145-26870-22135 | STM4312 | Hypothetical protein | Unknown - Regulatory target of HilD (1) | |

| | | | | | | | | | | | | |
|---|---|---|---|---|---|---|---|---|---|---|---|---|
| -2.30 | 0.52 | -0.54 | -1.38 | 1.75 | -1.90 | RS08675 | *yciA* | *yciA* | STM1736 | Acyl-CoA thioester hydrolase | Unknown | |
| -2.33 | -0.45 † | -0.58 | -2.53 | 1.75 | -2.08 | RS17000 | tRNA0057 | *secG-* tRNA0057 | | tRNA-Leu (gag) | tRNA & related | |
| -2.38 | -1.88 | -0.63 | -3.42 | 1.75 | -1.54 | RS19200 | *rpmG* | *rpmG* | STM3727 | 50S ribosomal protein L33 | Ribosome | ppGpp / DksA repressed |
| -2.33 | 1.35 | -0.60 | 0.17 † | 1.73 | -1.18 | RS19380 | *mgtC* | *mgtC* | STM3764 | Regulator of phosphate uptake | Phosphate uptake | |
| -2.38 | 0.31 † | -0.65 | -1.50 | 1.73 | -1.81 | RS00940 | *gluQRS* | *dksA-gluQRS* | STM0185 | tRNA glutamyl-Q(34) synthetase | tRNA & related | ppGpp / DksA repressed |
| -1.98 | -1.13 | -0.25 | -2.70 | 1.72 | -1.57 | RS16385 | *ygiR* | *ygiR* | STM3168 | YgiQ family radical SAM protein | Unknown | |
| -2.19 | -1.84 | -0.49 | -3.65 | 1.71 | -1.81 | RS10640 | *plaP* | *yoeI-plaP* | 0 | Putrescine/proton symporter | Polyamines - Putrescine import | |
| -2.38 | -1.91 | -0.68 | -2.59 | 1.70 | -0.68 | RS21335 | RS21335 | RS21335 | STM4156 | Hypothetical protein | Unknown | |
| -1.34 | -0.72 † | 0.36 | -2.04 | 1.70 | -1.32 | RS08885 | *prs* | *prs* | STM1780 | Ribose-phosphate pyrophosphokinase | Nucleotide metabolism - Pyrimidine biosynthesis | PurR repressed |
| -2.10 | -0.49 † | -0.44 | -2.02 | 1.66 | -1.53 | RS19185 | *waaA / kdtA* | *waaA-coaD* | STM3724 | 3-deoxy-D-manno-octulosonic acid transferase | Envelope - LPS synthesis | ArgR repressed |
| -2.15 | -0.59 † | -0.49 | -2.26 | 1.66 | -1.68 | RS12945 | *trmG/rlmN* | *trmG* | STM2525 | Bifunctional tRNA (adenosine(37)-C2)-methyltransferase TrmG/ribosomal RNA large subunit methyltransferase | tRNA & related | |
| -2.06 | -0.08 † | -0.40 | -2.18 | 1.65 | -2.10 | RS19040 | *trmL* | *trmL* | STM3695 | tRNA (uridine(34)/cytosine(34)/5-carboxymethylaminomethyluridine(34)-2'-O)-methyltransferase | tRNA & related | |
| -2.30 | -0.90 | -0.65 | -2.05 | 1.65 | -1.15 | RS07240 | *rsxC* | *rsxABCDGE-nth* | STM1457 | Electron transport complex subunit RsxC | ROS defence | |
| -2.43 | -0.56 † | -0.78 | -1.90 | 1.64 | -1.34 | RS07250 | *rsxA* | *rsxABCDGE-nth* | STM1459 | Electron transport complex subunit RsxA | ROS defence | |
| -2.40 | -0.38 † | -0.76 | -1.84 | 1.64 | -1.47 | RS02460 | *apt* | *apt* | STM0483 | Adenine phosphoribosyltransferase | Nucleotide metabolism - Purine salvage | GlaR repressed |
| -2.26 | -0.81 | -0.63 | -2.37 | 1.63 | -1.57 | RS10925 | *yegD* | *yegD* | STM2125 | Putative molecular chaperone | Unknown | |
| -2.02 | -1.75 | -0.40 | -3.09 | 1.62 | -1.33 | RS17485 | *fis* | *dusB-fis* | STM3385 | DNA-binding transcriptional regulator Fis | Regulation | IHF activated, ppGpp / DksA and Fis repressed |
| -2.24 | -0.29 † | -0.64 | -1.75 | 1.60 | -1.47 | RS26790 | *yqgC* | *yqgC* | STM3088 | Putative cytoplasmic protein | Unknown | |
| -1.98 | -0.90 † | -0.39 | -2.14 | 1.59 | -1.24 | RS07105 | *purR* | *purR* | STM1430 | HTH-type transcriptional repressor PurR | Nucleotide metabolism - Purine biosynthesis | PurR, Fur repressed |
| -1.30 | -1.11 | 0.28 | -2.25 | 1.58 | -1.14 | RS18685 | *yjhV* | *yjhV* | STM3625 | Putative transporter | Phage related | |
| -2.26 | -0.56 † | -0.69 | -2.33 | 1.58 | -1.77 | RS15365 | *sdaC* | *sdaCB* | STM2970 | HAAAP family serine/threonine permease | AA metabolism - Serine uptake | Lrp repressed |
| -1.91 | 1.00 | -0.34 | -0.77 | 1.57 | -1.77 | RS13225 | *rnc* | *rnc-era-recO-pdxJ-acpS* | STM2581 | ribonuclease III | Ribosome | |
| -1.17 | 1.64 | 0.38 | -0.12 † | 1.56 | -1.76 | RS14025 | *dprA* | *dprA* | STM3405 | DNA-processing protein DprA | Transformation | |
| -2.06 | 0.07 † | -0.52 | -1.54 | 1.55 | -1.61 | RS02175 | *thiI* | *thiI* | STM0425 | tRNA 4-thiouridine(8) synthase ThiI | tRNA & related | SAM repressed |
| -1.86 | 0.04 † | -0.32 | -1.67 | 1.54 | -1.71 | RS16645 | *rlmG* | *rlmG* | STM3220 | 23S rRNA (guanine(1835)-N(2))-methyltransferase RlmG | Ribosome | |
| -2.15 | -0.82 | -0.62 | -1.78 | 1.53 | -0.96 | RS07235 | *rsxD* | *rsxABCDGE-nth* | STM1456 | Electron transport complex subunit RsxD | ROS defence | |
| -1.87 | -1.40 | -0.34 | -3.02 | 1.52 | -1.62 | RS21275 | tRNA0076 | tRNA0076-77 | | tRNA-Gly (tcc) | tRNA & related | |
| -2.29 | -2.07 | -0.77 | -2.25 | 1.52 | -0.17 | RS22145 | RS22145 | RS22150-22145-26870-22135 | STM4314 | GerE family regulatory protein | Unknown - Operon is regulatory target of HilD (1) | |

| | | | | | | | | | | | | |
|---|---|---|---|---|---|---|---|---|---|---|---|---|
| -1.81 | -1.83 | -0.32 | -3.06 | 1.50 | -1.23 | RS18890 | *avtA* | *avtA* | STM3665 | Valine - pyruvate transaminase | AA metabolism - Alanine biosynthesis | Lrp regulated |
| -1.84 | -1.32 † | -0.37 | -2.72 † | 1.46 | -1.41 | RS03170 | *pagP* | *pagP* | STM0628 | Lipid IV(A) palmitoyltransferase PagP | Envelope - Lipid A biosynthesis - PhoP activated in Salmonella (2) | PhoP, SlyA activated, H-NS repressed |
| -2.19 | -2.32 | -0.72 | -3.52 | 1.46 | -1.21 | RS00840 | *speD* | *speED* | STM0165 | Adenosylmethionine decarboxylase | Polyamines - Spermidine synthesis | |
| -2.11 | -0.37 † | -0.66 | -1.48 | 1.45 | -1.11 | RS21255 | *coaA/panK* | *coaA* | STM4139 | Type I pantothenate kinase | CoA synthesis | |
| -1.89 | 2.92 | -0.44 | -0.70 † | 1.45 | -3.62 | RS12265 | *fadL* | *fadL* | STM2391 | Long-chain fatty acid transporter FadL | Fatty acid uptake | Lrp, PdhR, CRP, PhoP, ppGpp activated, RpoN, ArcA, FadR, OmpR, Rob repressed. |
| -1.71 | -2.21 | -0.27 | -3.25 | 1.44 | -1.04 | RS19780 | *yidC* | *rpmHA-yidDC* | STM3842 | Membrane protein insertase YidC | Envelope – IM protein insertion | |
| -1.81 | -0.35 † | -0.38 | -1.69 | 1.42 | -1.34 | RS08595 | *rluB* | *rluB* | STM1719 | 23S rRNA pseudouridine(2605) synthase RluB | Ribosome | |
| -1.72 | -0.45 † | -0.31 | -1.73 | 1.41 | -1.28 | RS08855 | *sirB2* | *hemA-prfA-prmC-sirB2B1-kdsA* | STM1774 | SirB family protein | Linked to SPI-1? | |
| -2.15 | -0.94 † | -0.75 | -2.63 | 1.41 | -1.69 | RS15370 | *sdaB* | *sdaCB* | STM2971 | L-serine ammonia-lyase | AA metabolism - Serine degradation | |
| -1.96 | -2.10 | -0.56 | -3.01 † | 1.40 | -0.91 | RS13800 | *rpsP* | *rpsP* | STM2676 | 30S ribosomal protein S16 | Ribosome | |
| -1.14 | 0.71 | 0.26 | -0.47 | 1.39 | -1.17 | RS23350 | RS23350 | RS23350 | STM4551 | GGDEF domain-containing protein | Signalling? | |
| -2.60 | -1.02 | -1.24 | -2.15 | 1.36 | -1.12 | RS04770 | *rpsA* | *rpsA* | STM0981 | 30S ribosomal protein S1 | Ribosome | ppGpp / DksA repressed |
| -1.62 | -1.60 | -0.27 | -2.98 | 1.35 | -1.37 | RS17800 | *rpsL* | *rpsL* | STM3448 | 30S ribosomal protein S12 | Ribosome | ppGpp / DksA repressed |
| -1.90 | -0.67 | -0.55 | -2.48 | 1.34 | -1.81 | RS00045 | *satP* | *satP* | STM0009 | Acetate / succinate symporter | Acetate homeostasis | |
| -1.86 | -2.47 | -0.54 | -3.43 | 1.32 | -0.96 | RS22540 | *rpsF* | *rpsF-priB-rpsR* | STM4391 | 30S ribosomal protein S6 | Ribosome | ppGpp repressed |
| -2.10 | -1.47 | -0.79 | -2.73 | 1.32 | -1.25 | RS16955 | *rpsO* | *rpsO* | STM3283 | 30S ribosomal protein S15 | Ribosome | ppGpp repressed |
| -2.04 | 1.86 | -0.73 | 0.04 † | 1.32 | -1.82 | RS12245 | *sixA* | *sixA* | STM2387 | Phosphohistidine phosphatase SixA | Signalling | |
| -2.36 | 1.01 † | -1.05 | -1.19 | 1.31 | -2.20 | RS12950 | *ndk* | *ndk* | STM2526 | Nucleoside-diphosphate kinase | Nucleotide metabolism - salvage / biosynthesis | ArcA repressed |
| -2.00 | 2.18 | -0.70 | 0.47 † | 1.30 | -1.70 | RS08955 | *cbdX* | *appCB-cbdX* | STM1794 | Cytochrome bd-II oxidase subunit CbdX | Respiration | AppY, ArcA, YdeO activated |
| -3.41 | -1.21 | -2.12 | -1.31 | 1.29 | -0.10 | RS14945 | *spaS* | SPI-1 | STM2887 | SPI-1 type III secretion system export apparatus protein SpaS | SPI-1 | |
| -1.73 | -2.62 | -0.45 | -3.47 | 1.28 | -0.85 | RS22550 | *rpsR* | *rpsF-priB-rpsR* | STM4393 | 30S ribosomal protein S18 | Ribosome | ppGpp repressed |
| -1.84 | -0.66 † | -0.56 | -1.96 | 1.27 | -1.29 | RS02715 | *lpxH* | *ppiB- lpxH* | STM0535 | UDP-2,3-diacylglucosamine diphosphatase | Envelope - Lipid A synthesis | |
| -2.42 | 0.02 † | -1.15 | -0.90 | 1.27 | -0.92 | RS06140 | *envE* | *envE* | STM1242 | Lipoprotein | SPI-11 | |
| -2.40 | -1.25 | -1.14 | -1.91 | 1.27 | -0.66 | RS14855 | *orgB* | SPI-1 | STM2868 | Oxygen-regulated invasion protein | SPI-1 | |
| -1.81 | -0.69 | -0.55 | -1.78 | 1.26 | -1.09 | RS07245 | *rsxB* | *rsxABCDGE-nth* | STM1458 | Electron transport complex subunit RsxB | ROS defence | |
| -1.77 | -0.43 † | -0.53 | -1.97 | 1.25 | -1.54 | RS18495 | *pitA* | *pitA* | STM3589 | Inorganic phosphate transporter PitA | Phosphate uptake | FNR activated |
| -1.65 | -1.93 | -0.41 | -3.14 | 1.24 | -1.20 | RS00845 | *speE* | *speED* | STM0166 | Polyamine aminopropyltransferase | Polyamines - Spermidine synthesis | |
| -1.69 | -0.12 † | -0.46 | -1.81 | 1.24 | -1.70 | RS27005 | *yoaK* | *yoaK* | | YoaK family small membrane protein | Unknown | |
| -1.78 | -2.81 | -0.56 | -3.58 | 1.23 | -0.77 | RS22545 | *priB* | *rpsF-priB-rpsR* | STM4392 | Primosomal replication protein N | DNA replication | ppGpp repressed in coli |

| | | | | | | | | | | | | |
|---|---|---|---|---|---|---|---|---|---|---|---|---|
| -2.19 | 0.04 † | -0.96 | -1.08 | 1.23 | -1.13 | RS01120 | *ispU* | *ispU-cdsA-resP-bamA* | STM0221 | (2E,6E)-farnesyl-diphosphate-specific ditrans,polycis-undecaprenyl-diphosphate synthase | UPP synthesis | |
| -2.05 | -2.33 | -0.83 | -1.79 | 1.22 | 0.55 | RS26870 | RS26870 | RS22150-22145-26870-22135 | STM4313 | Hypothetical protein | Unknown - Regulatory target of HilD (1) | |
| -1.64 | -0.03 † | -0.42 | -0.91 | 1.22 | -0.88 | RS23295 | RS23295 | RS23270-95 | STM4540 | SIS domain-containing protein | PTS system? | |
| -2.14 | -0.16 † | -0.92 | -1.36 | 1.22 | -1.20 | RS19935 | *atpI* | *atpIBEFHAGDC* | STM3872 | F0F1 ATP synthase subunit I | Energy metabolism | |
| -1.95 | -0.48 † | -0.73 | -1.34 | 1.22 | -0.86 | RS08500 | *rnb* | *rnb* | STM1702 | Exoribonuclease II | RNA processing | |
| -3.31 | -1.23 | -2.10 | -1.43 | 1.22 | -0.20 | RS14950 | *spaR* | SPI-1 | STM2888 | SPI-1 type III secretion system export apparatus protein SpaR | SPI-1 | |
| -1.50 | 1.01 | -0.29 | -0.39 † | 1.21 | -1.39 | RS08250 | RS08250 | RS08250-55 | | Pseudogene | Unknown | |
| -1.52 | -0.01 † | -0.31 | -1.31 | 1.21 | -1.30 | RS07125 | *rnt* | *rnt* | STM1434 | Ribonuclease T | RNA processing | |
| -4.02 | -0.03 † | -2.82 | -1.46 | 1.20 | -1.43 | RS22130 | RS22130 | RS22130 | STM4310 | YjiK family protein/ putative inner membrane protein | Unknown - Regulatory target of HilD (1) | |
| -2.90 | -1.08 | -1.72 | -0.96 | 1.18 | 0.12 | RS14975 | *spaM* | SPI-1 | STM2893 | SPI-1 type III secretion system protein | SPI-1 | |
| -1.45 | -0.79 | -0.26 | -1.65 | 1.18 | -0.86 | RS17420 | *mreC* | *mreBCD*-RS17410-*rng* | STM3373 | Rod shape-determining protein | Cell division / shape control | Nac, BolA repressed |
| -1.95 | -2.47 | -0.77 | -3.02 | 1.18 | -0.55 | RS13795 | *rimM* | *rpsP-rimM-trmD-rplS* | STM2675 | Ribosome maturation factor | Ribosome | ppGpp / DksA, FNR repressed |
| -1.94 | 1.53 | -0.77 | 0.09 † | 1.17 | -1.44 | RS04440 | *potF* | *potFGHI* | STM0877 | Spermidine/putrescine ABC transporter substrate binding protein | Polyamines - Putrescine import | ArgR repressed, Lrp regulated, NtrC activated ArcA regulated |
| -1.81 | -0.61 | -0.65 | -1.72 | 1.16 | -1.10 | RS09220 | *proQ* | *proQ-prc* | STM1846 | RNA chaperone ProQ | RNA processing | |
| -1.36 | -2.19 | -0.20 | -2.95 | 1.16 | -0.76 | RS11425 | *rplY* | *rplY* | STM2224 | 50S ribosomal protein L25 | Ribosome | ppGpp / DksA repressed |
| -2.09 | -1.21 | -0.93 | -1.94 | 1.15 | -0.73 | RS15975 | *yqgB* | *yqgB-speA* | STM3087 | Acid stress response protein | Acid stress | |
| -1.56 | -0.39 † | -0.41 | -1.64 | 1.15 | -1.25 | RS23380 | *holD* | *holD-rimI-yjjG* | STM4557 | DNA polymerase III subunit ψ | DNA replication | |
| -1.82 | -0.42 † | -0.67 | -1.65 | 1.15 | -1.23 | RS22410 | *yjeT* | *yjeT* | STM4365 | DUF2065 domain-containing protein | Unknown | |
| -1.68 | -2.39 | -0.54 | -3.01 | 1.14 | -0.62 | RS17280 | *rplM* | *rplM-rpsI* | STM3345 | 50S ribosomal protein L13 | Ribosome | ppGpp / DksA repressed |
| -1.80 | -2.40 | -0.66 | -3.27 | 1.14 | -0.87 | RS22555 | *rplI* | *rpsF-priB-rpsR-rplI* | STM4394 | 50S ribosomal protein L9 | Ribosome | CRP activated, ppGpp repressed |
| -1.57 | -1.04 | -0.43 | -1.93 | 1.14 | -0.90 | RS05880 | *plsX* | *plsX-fabHDG* | STM1192 | Phosphate acyltransferase PlsX | Fatty acid metabolism? | |
| -1.35 | -0.35 † | -0.22 | -1.11 | 1.13 | -0.77 | RS19285 | *recG* | *gmk-rpoZ-spoT-trmH-recG* | STM3744 | ATP-dependent DNA helicase RecG | DNA repair | CreB activated, ppGpp / DksA repressed |
| -1.57 | -0.97 | -0.44 | -1.75 | 1.13 | -0.78 | RS04890 | *pcnB* | *pcnB-folK* | STM0184 | Nicotinate phosphoribosyltransferase | Plasmid copy number | ppGpp / DksA repressed |
| -2.32 | -1.04 | -1.20 | -2.06 | 1.12 | -1.02 | RS04150 | *rhlE* | *rhlE* | STM0820 | ATP-dependent RNA helicase RhlE | Ribosome | CecR activated |
| -1.37 | -0.89 | -0.24 | -1.49 | 1.12 | -0.60 | RS20200 | *wecF* | *rfe-wzzE-wecBC-rffGHC-wecE-wzxE-wecF-wzyE-rffM-yifK* | STM3927 | TDP-N-acetylfucosamine:lipid II N-acetylfucosaminyltransferase | Envelope - ECA synthesis | NsrR repressed |
| -2.31 | -2.46 | -1.19 | -2.34 | 1.11 | 0.12 | RS14845 | *sirC* | SPI-1 | STM2867 | Transcriptional regulator | SPI-1 | |
| -1.61 | -1.00 | -0.50 | -1.80 | 1.11 | -0.80 | RS22535 | RS22535 | RS22535 | STM4390 | Hypothetical protein | Unknown | |
| -3.60 | -0.95 | -2.49 | -1.70 | 1.11 | -0.75 | RS09310 | RS09310 | RS09310-5 | STM1863 | Putative DUF5993 family protein | Unknown | |

| | | | | | | | | | | | | |
|---|---|---|---|---|---|---|---|---|---|---|---|---|
| -1.58 | 0.40 † | -0.47 | -0.85 | 1.11 | -1.25 | RS21040 | *priA* | *priA* | STM4095 | primosomal protein N | DNA replication | |
| -1.61 | -2.50 | -0.50 | -3.14 | 1.10 | -0.64 | RS17760 | *rplC* | *rpsJ-rplCDWB-rpsS-rplV-rpsC-rplP-rpmC-rpsQ* | STM3440 | 50S ribosomal protein L3 | Ribosome | ppGpp / DksA repressed |
| -1.30 | 0.16 † | -0.21 | -1.00 | 1.08 | -1.15 | RS13220 | *era* | *rnc-era-recO-pdxJ-acpS* | STM2580 | GTPase Era | Ribosome | |
| -1.47 | 0.03 † | -0.39 | -1.00 | 1.08 | -1.03 | RS11145 | *pbpG* | *pbpG* | STM2168 | D-alanyl-D-alanine endopeptidase (PBP7) | Envelope - Peptidoglycan synthesis / modification | |
| -1.52 | -2.38 | -0.44 | -3.26 | 1.08 | -0.87 | RS17645 | *rpsK* | *rpsMKD-rpoA* | STM3417 | 30S ribosomal protein S11 | Ribosome | ppGpp / DksA repressed in coli |
| -1.34 | -0.44 † | -0.26 | -1.38 | 1.08 | -0.94 | RS18025 | *mrcA* | *mrcA* | STM3493 | Peptidoglycan glycosyltransferase/peptidoglycan DD-transpeptidase (PBP1a) | Envelope - Peptidoglycan synthesis / modification | |
| -1.73 | 1.21 | -0.65 | 0.45 | 1.08 | -0.76 | RS04350 | RS04350 | RS04350 | STM0859 | Putative LysR family transcriptional regulator | Regulation | |
| -1.39 | 0.82 | -0.31 | -0.72 | 1.08 | -1.53 | RS13620 | *pssA* | *pssA* | STM2652 | CDP-diacylglycerol--serine O-phosphatidyltransferase | Envelope - Phospholipid synthesis | |
| -1.41 | -1.00 | -0.34 | -1.87 | 1.07 | -0.86 | RS16980 | *rimP* | *rimP-nusA-infB* | STM3288 | Ribosome maturation factor | Ribosome | Fis activated, Nac, ArgR, CRP repressed |
| -1.52 | -0.08 † | -0.46 | -1.23 | 1.07 | -1.15 | RS08255 | *ttcA* | RS08250-*ttcA* | STM1654 | tRNA 2-thiocytidine(32) synthetase | tRNA & related | |
| -1.66 | -0.11 † | -0.60 | -1.51 | 1.06 | -1.40 | RS15635 | RS15635 | RS15635 | STM3022 | Putative amino acid permease | AA metabolism - Amino acid transport | |
| -1.73 | -1.96 | -0.68 | -2.11 | 1.05 | -0.15 | RS22150 | RS22150 | RS22150-22145-26870-22135 | STM4315 | AraC family transcriptional regulator | Unknown - Operon is regulatory target of HilD (1) | |
| -1.68 | 0.35 | -0.64 | -0.77 | 1.04 | -1.12 | RS00445 | *folA* | *folA* | STM0087 | Type 3 dihydrofolate reductase | Tetrahydrofolate biosynthesis | IHF, TyrR activated |
| -3.22 | 1.02 | -2.18 | -0.20 † | 1.04 | -1.22 | RS03005 | *entC* | *entCEBAH* | STM0595 | Isochorismate synthase | Iron acquisition | CRP activated, Fur repressed |
| -2.22 | -1.72 | -1.18 | -1.91 | 1.04 | -0.20 | RS14850 | *orgC* | SPI-1 | STM2868 | Type III secretion system effector protein | SPI-1 | |
| -2.48 | -1.48 | -1.45 | -1.64 | 1.03 | -0.16 | RS14985 | *spaK* | SPI-1 | STM2895 | SPI-1 type III secretion system chaperone | SPI-1 | |
| -1.57 | 0.30 † | -0.54 | -0.90 | 1.03 | -1.20 | RS06175 | tRNA0022 | tRNA0022 | 0 | tRNA-Arg | tRNA & related | |
| -1.32 | 0.08 † | -0.29 | -0.97 | 1.03 | -1.05 | RS20445 | *rfaH* | *rfaH* | STM3977 | Transcription/translation regulatory transformer protein | Envelope – LPS synthesis | Nac activated |
| -1.53 | -2.01 | -0.50 | -2.73 | 1.03 | -0.72 | RS17710 | *rplN* | *rplNXE-rpsNH-rplFR-rpsE-rpmD-rp10-secY-rpmJ* | STM3430 | 50S ribosomal protein L14 | Ribosome | ppGpp / DksA repressed in coli |
| -2.42 | 2.23 | -1.39 | 1.46 | 1.03 | -0.77 | RS23105 | RS23105 | RS23105 | STM4504 | Hypothetical protein | Unknown | |
| -1.75 | 4.70 | -0.73 | 3.20 | 1.02 | -1.51 | RS21000 | *glpF* | *glpFKX* | STM4087 | Glycerol facilitator | Glycerol uptake | GRP activated, GlpR repressed |
| -1.77 | -0.18 † | -0.76 | -1.25 | 1.02 | -1.06 | RS17035 | *yhbY* | *yhbY* | STM3298 | Ribosome assembly factor | Ribosome | Lrp repressed |
| -1.30 | -0.94 | -0.28 | -1.57 | 1.02 | -0.63 | RS01160 | *lpxB* | *bamA-skp-lpxD-fabZ-lpxAB-rnhA-dnaE-accA* | STM0229 | Lipid A disaccharide synthase | Envelope – LPS synthesis | $\sigma^{24}$ activated |
| -1.24 | 0.20 † | -0.23 | -1.05 | 1.02 | -1.25 | RS13575 | *srmB* | *srmB* | STM2643 | ATP-dependent RNA helicase | Ribosome | |
| -1.58 | 0.94 | -0.57 | -0.03 † | 1.01 | -0.97 | RS05475 | *scsB* | RS05470-80-*agp* | STM1114 | Protein-disulphide reductase | Disulphide bond formation | |
| -1.30 | -1.47 | -0.30 | -2.41 | 1.00 | -0.94 | RS05245 | *lonH* | *lonH* | STM1068 | Lon protease family protein | | |

**Key to columns (left to right)**

**$Log_2$ fold change** values for each gene: 1 h vs 3 h in the wild type; 3 h vs 5 h in the wild type; 1 h vs 3 h in Δ*acrB*; 3 h vs 5 h in Δ*acrB*.

**ΔLog$_2$ values** for the 1 h vs 3 h and 3 h vs 5 h comparisons. Δlog$_2$ values are calculated by Log$_2$ fold change for Δ*acrB* minus Log$_2$ fold change for the wild type. Positive values indicate more upregulation or less downregulation in wt, negative values more upregulation or less downregulation in *acrB*. Genes are sorted in decreasing Δlog$_2$ 1vs3 h order.

**ID**: locus name.

**Gene name**

**Operon structure:** if unknown are inferred from genomic sequence and comparison with ecocyc.com.

**STM**: STM gene number.

**Product**: Function of gene product, if known.

**Comments**: General classification of function of gene product.

**Regulation in *E. coli***: sourced from Ecocyc.com unless otherwise referenced.

**Colour mapping:**

| | | | | | | | | |
|---|---|---|---|---|---|---|---|---|
| -4 | -3 | -2 | -1 | 0 | 1 | 2 | 3 | 4 |

Log$_2$ fold change values that are non-significant ($p_{adj} > 0.05$) are in white and marked with a dagger †. ΔLog$_2$ fold change values derived from one or more non-significant Log$_2$ fold change values are also in white.

### Supplemental Table S2. Genes in group 2 - Genes that are more upregulated at 3hr in Δ*acrB* than wt – “early activation” pattern

| Log$_2$ fold change | | | | *acrB* - wt Δlog$_2$ | | | | | | | | | | |
|---|---|---|---|---|---|---|---|---|---|---|---|---|---|---|
| wt 1vs3 | wt 3vs5 | *acrB* 1vs3 | *acrB* 3vs5 | 1vs3 | 3vs5 | ID | Gene name | Operon | STM | Product | Comments | Regulation in *Salmonella* | Regulation in *E. coli* | SalComMac log$_2$ fold change Anaerobic shock |
| 0.84 | 3.50 | 3.85 | 0.08 † | 3.01 | -3.41 | SL1344_RS05365 | *orfX* | *orfX* | STM1092 | Hypothetical YlcI/YnfO family protein | Part of SPI-5 | DksA repressed (3) | | 5.23 |
| 2.12 | -1.18 | 4.32 | -2.21 | 2.20 | -1.03 | SL1344_RS06235 | *yntB / nikB* | *yntABCDE / nikABCDE* | STM1256 | ABC transporter permease | High-affinity nickel transporter | | FNR activated, NikR repressed | 2.25 |
| 1.02 | -0.79 † | 3.18 | -2.17 | 2.15 | -1.38 | SL1344_RS06240 | *yntC / nikC* | *yntABCDE / nikABCDE* | STM1257 | ABC transporter permease | High-affinity nickel transporter | | FNR activated, NikR repressed | 1.93 |
| 0.82 | -0.75 | 2.93 | -1.81 | 2.11 | -1.05 | SL1344_RS06250 | *yntE / nikE* | *yntABCDE / nikABCDE* | STM1259 | Peptide ABC transporter ATP-binding protein | High-affinity nickel transporter | | FNR activated, NikR repressed | 1.68 |
| 3.40 | -2.06 | 5.40 | -3.19 | 2.00 | -1.14 | SL1344_RS06230 | *yntA / nikA* | *yntABCDE / nikABCDE* | STM1255 | Nickel ABC transporter substrate-binding protein | High-affinity nickel transporter | | FNR activated, NikR repressed | 4.09 |
| -0.62 | -1.78 | 1.29 | -3.26 | 1.91 | -1.47 | SL1344_RS19800 | | 19800- 19805 | STM3845 | Hypothetical protein (retron St85 family effector protein) | Anti-phage defence | | | -1.25 |
| -0.66 | 1.24 | 1.03 | -0.31 † | 1.69 | -1.56 | SL1344_RS16765 | *tdcB* | *tdcABCD-pflB-tdcG* | STM3244 | Bifunctional threonine ammonia-lyase/L-serine ammonia-lyase | Anaerobic serine / threonine degradation | H-NS repressed, TdcA activated (4) | FNR, CRP, IHF, TdcA, TdcR activated | 7.45 |
| -0.89 | 0.64 † | 0.79 | -0.72 | 1.68 | -1.35 | SL1344_RS09910 | *fliR* | *fliLMNOPQR* | STM1981 | Flagellar type III secretion system protein | Flagella - Class 2 flagellar gene | | $\sigma^{70}$/ $\sigma^{28}$, FlhDC activated | -2.12 |
| 1.79 | 0.27 † | 3.46 | -1.91 | 1.67 | -2.18 | SL1344_RS15890 | | 15890-15910 | STM3071 | Putative DNA-binding protein | Putative $Co^{2+}$ / $Ni^{2+}$ transporter | | | 2.14 |
| -0.49 | 0.23 † | 1.14 | -2.11 | 1.62 | -2.34 | SL1344_RS09690 | *tyrP* | *tyrP* | STM1937 | Tyrosine transporter TyrP | Tyrosine import | | TyrR, Lrp regulated, IHF repressed | -1.51 |
| 1.49 | -0.53 † | 3.04 | -2.28 | 1.55 | -1.75 | SL1344_RS11590 | *napB* | *napFDAGHBC-ccmABCDEFG2* | STM2256 | nitrate reductase cytochrome c-type subunit | Anaerobic respiration – Periplasmic nitrate reductase | | FNR, ModE,FlhCD activated, MArP regulated, NarL, IscR repressed | 4.69 |
| -0.77 | -0.19 † | 0.77 | -1.41 | 1.53 | -1.22 | SL1344_RS14220 | | 14220 | STM2747 | DUF4435 domain-containing protein | Unknown function | | | -0.18 |
| 2.45 | -1.51 | 3.95 | -2.30 | 1.50 | -0.79 | SL1344_RS06245 | *yntD / nikD* | *yntABCDE / nikABCDE* | STM1258 | ATP-binding cassette domain-containing protein | High-affinity nickel transporter | | FNR activated, NikR repressed | 1.74 |
| 2.11 | -0.63 † | 3.59 | -2.68 | 1.48 | -2.05 | SL1344_RS11600 | *napG* | *napFDAGHBC-ccmABCDEFG2* | STM2258 | ferredoxin-type protein NapG | Anaerobic respiration – Periplasmic nitrate reductase | | FNR, ModE,FlhCD activated, NarP regulated, NarL, IscR repressed | 4.78 |
| -0.66 | 3.84 | 0.81 | 2.43 | 1.46 | -1.41 | SL1344_RS11710 | | 11710 | STM2281 | LysR family transcriptional regulator | Unknown function | | | 1.92 |
| 1.81 | -1.12 | 3.26 | -2.37 | 1.45 | -1.25 | SL1344_RS11560 | *ccmE2* | *napFDAGHBC-ccmABCDEFG2* | STM2250 | cytochrome c maturation protein CcmE | Cytochrome *c* maturation – cluster 2 | | FNR, ModE, FlhCD activated, NarP regulated, NarL, IscR repressed | 0.32 |
| 2.60 | 0.96 | 4.05 | -0.25 † | 1.45 | -1.21 | SL1344_RS19665 | *yhjA / ccp* | *ccp-?-ccmABSCDEFG1* | STM3820 | c-type cytochrome | Cytochrome *c* peroxidase | FNR activated(5) | FNR, OxyR activated | 0.03 |
| -0.93 | 3.58 | 0.41 | 2.21 | 1.34 | -1.38 | SL1344_RS19740 | | 19740 | STM3834 | LysR family transcriptional regulator | Unknown function | | | 3.13 |
| -0.85 | -0.53 † | 0.48 | -2.18 | 1.33 | -1.64 | SL1344_RS15045 | *pphB* | *pphB* | STM2907 | Serine/threonine protein phosphatase | SPI-1 | | Nac repressed | -0.79 |
| 1.89 | -0.97 † | 3.21 | -2.68 | 1.32 | -1.71 | SL1344_RS11605 | *napA* | *napFDAGHBC-ccmABCDEFG2* | STM2259 | nitrate reductase catalytic subunit NapA | Anaerobic respiration – Periplasmic nitrate reductase | | FNR, ModE,FlhCD activated, NarP regulated, NarL, IscR repressed | 5.38 |
| 2.13 | -1.39 | 3.45 | -2.46 | 1.32 | -1.07 | SL1344_RS11555 | *ccmF2* | *napFDAGHBC-ccmABCDEFG2* | STM2249 | c-type cytochrome biogenesis protein CcmF | Cytochrome *c* maturation – cluster 2 | | FNR, ModE,FlhCD activated, NarP regulated, NarL, IscR repressed | 0.00 |
| 0.73 | -3.01 | 2.04 | -3.07 | 1.32 | -0.06 | SL1344_RS09850 | *fliF* | *fliFGHIJKL* | STM1969 | flagellar M-ring protein FliF | Flagella - Class 2 flagellar gene | FNR, FlhDC activated (5, 6) | $\sigma^{70}$/ $\sigma^{28}$, FlhDC activated, CsgD repressed | -2.64 |
| 2.00 | 0.50 † | 3.31 | -1.31 | 1.32 | -1.81 | SL1344_RS21950 | *nrfA* | *nrfABCDEFG* | STM4277 | ammonia-forming nitrite reductase cytochrome c552 subunit | Anaerobic respiration – Periplasmic nitrite reductase | FNR activated (5) | FNR,FlhDC, NarP activated, IHF, NarL regulated, Fis, NsrR repressed | 5.42 |
| -0.58 | 0.18 † | 0.69 | -0.88 | 1.27 | -1.06 | SL1344_RS03870 | | 03870 | STM0764 | LysR family transcriptional regulator | Unknown function | | | 0.00 |

| | | | | | | | | | | | | | |
|---|---|---|---|---|---|---|---|---|---|---|---|---|---|
| 0.70 | -0.73 † | 1.95 | -1.83 | 1.25 | -1.10 | SL1344_RS09685 | *yecH* | *yecH* | STM1936 | Hypothetical protein | Unknown function – divergent from *tyrP* | Downregulated by adrenaline (7) | | 2.74 |
| -0.69 | -0.47 | 0.55 | -1.48 | 1.24 | -1.01 | SL1344_RS11335 | *setB* | *setB* | STM2207 | Sugar efflux transporter SetB | Sugar efflux | | | 0.00 |
| -0.91 | 0.37 † | 0.33 | -0.58 | 1.24 | -0.95 | SL1344_RS13240 | *gogB* | *gogB* | STM2584 | Phage-encoded type III secretion effector GogB | Phage | | | -0.29 |
| 0.80 | -1.19 | 2.03 | -1.89 | 1.23 | -0.70 | SL1344_RS07940 | *srfA* | *srfABC* | STM1593 | SsrAB-activated protein | Proposed virulence effector; class 2 flagellar gene | FNR, FlhDC activated (5, 6) | | -2.47 |
| 1.34 | -1.64 | 2.56 | -2.57 | 1.22 | -0.93 | SL1344_RS11545 | *ccmH2* | *napFDAGHBC-ccmABCDEFG2* | STM2247 | cytochrome c-type biogenesis protein CcmH | Cytochrome *c* maturation – cluster 2 | | FNR, ModE,FlhCD activated, NarP regulated, NarL, IscR repressed | 2.41 |
| 2.05 | -1.55 | 3.26 | -2.08 | 1.22 | -0.52 | SL1344_RS11565 | *ccmD2* | *napFDAGHBC-ccmABCDEFG2* | STM2251 | heme exporter protein CcmD | Cytochrome *c* maturation – cluster 2 | | FNR, ModE,FlhCD activated, NarP regulated, NarL, IscR repressed | 0.00 |
| 1.86 | -0.72 † | 3.06 | -1.93 | 1.20 | -1.20 | SL1344_RS19660 | *ccmA1* | *ccp-?-ccmABSCDEFGH1* | STM3819 | cytochrome c biogenesis heme-transporting ATPase CcmA | Cytochrome *c* maturation – cluster 1 | | | |
| 1.56 | -0.63 † | 2.76 | -2.02 | 1.20 | -1.39 | SL1344_RS19655 | *ccmB1* | *ccp-?-ccmABSCDEFGH1* | STM3818 | heme exporter protein CcmB | Cytochrome *c* maturation – cluster 1 | | | 0.00 |
| 1.65 | -0.58 † | 2.85 | -2.28 | 1.20 | -1.70 | SL1344_RS11575 | *ccmB2* | *napFDAGHBC-ccmABCDEFG2* | STM2253 | heme exporter protein CcmB | Cytochrome *c* maturation – cluster 2 | | FNR, ModE,FlhCD activated, NarP regulated, NarL, IscR repressed | 0.00 |
| -0.47 | -0.41 † | 0.72 | -1.18 | 1.19 | -0.77 | SL1344_RS23215 | *hsdS* | *hsdMS* | STM4524 | restriction endonuclease subunit S | Restriction endonuclease | | | 0.26 |
| 2.26 | 2.32 | 3.43 | 0.80 † | 1.17 | -1.52 | SL1344_RS22105 | | 22105-22120 | STM4305 | molybdopterin-dependent oxidoreductase | Putative DMSO reductase | FNR activated (5) | | 4.01 |
| -0.74 | 5.35 | 0.43 | 3.70 | 1.17 | -1.65 | SL1344_RS20930 | *lsrR* | *lsrRK* | STM4073 | transcriptional regulator | Quorum sensing | | CRP activated, LsrR repressed | -0.36 |
| 2.03 | -0.72 † | 3.19 | -2.03 | 1.16 | -1.31 | SL1344_RS11580 | *ccmA2* | *napFDAGHBC-ccmABCDEFG2* | STM2254 | cytochrome c biogenesis heme-transporting ATPase CcmA | Cytochrome *c* maturation – cluster 2 | | FNR, ModE,FlhCD activated, NarP regulated, NarL, IscR repressed | 0.00 |
| 3.01 | -0.53 † | 4.15 | -1.78 | 1.14 | -1.24 | SL1344_RS16910 | *yhbU / ubiU* | *ubiUV* | STM3274 | $O_2$-independent ubiquinone biosynthesis protein | Ubiquinone biosynthesis | FNR activated (5) | Nac repressed | 3.57 |
| 1.86 | -0.23 † | 2.99 | -1.50 | 1.13 | -1.27 | SL1344_RS15900 | *ecfT* | 15890-15910 | STM3073 | Energy-coupling factor transporter transmembrane protein EcfT - cobalt transporter? | Putative $Co^{2+}$ / $Ni^{2+}$ transporter | | | 1.29 |
| -0.86 | -0.35 † | 0.26 | -1.40 | 1.12 | -1.05 | SL1344_RS08880 | *ispE / ipk* | *lolB-ispE* | STM1779 | 4-(cytidine 5'-diphospho)-2-C-methyl-D-erythritol kinase | Isoprenoid synthesis pathway | | | -1.64 |
| 0.74 | 4.78 | 1.85 | 3.40 † | 1.10 | -1.38 | SL1344_RS12010 | | 12020-12005 | STM2341 | Putative transketolase | Unknown function | FNR repressed (5) | | 4.45 |
| -0.48 | 3.56 | 0.61 | 2.48 | 1.09 | -1.08 | SL1344_RS21085 | | 21085 | STM4103 | Hypothetical protein | Unknown function | | | 1.63 |
| 2.14 | -1.46 | 3.21 | -2.52 | 1.07 | -1.06 | SL1344_RS19635 | *ccmF1* | *ccp-?-ccmABSCDEFGH1* | STM3814 | *c*-type cytochrome biogenesis protein CcmF | Cytochrome *c* maturation – cluster 1 | | | 0.00 |
| 1.99 | -0.82 | 3.04 | -1.90 | 1.05 | -1.08 | SL1344_RS19640 | *ccmE1* | *ccp-?-ccmABSCDEFGH1* | STM3815 | Cytochrome c maturation protein CcmE | Cytochrome *c* maturation – cluster 1 | | | 0.32 |
| 1.01 | -3.72 | 2.03 | -2.85 | 1.02 | 0.86 | SL1344_RS09860 | *fliH* | *fliFGHIJKL* | STM1971 | Flagellar assembly protein FliH | Flagella - Class 2 flagellar gene | FNR, FlhDC activated (5, 6) | $\sigma^{70}$/ $\sigma^{28}$, FlhDC activated, CsgD repressed | -2.56 |
| 1.79 | -1.27 | 2.79 | -2.88 | 1.00 | -1.61 | SL1344_RS11550 | *ccmG2* | *napFDAGHBC-ccmABCDEFG2* | STM2248 | Thiol:disulfide interchange protein | Cytochrome *c* maturation – cluster 2 | | FNR, ModE,FlhCD activated, NarP regulated, NarL, IscR repressed | 0.00 |

**Key to columns (left to right)**

**$Log_2$ fold change** values for each gene: 1 h vs 3 h in the wild type; 3 h vs 5 h in the wild type; 1 h vs 3 h in Δ*acrB*; 3 h vs 5 h in Δ*acrB*.

**$\Delta Log_2$ values** for the 1 h vs 3 h and 3 h vs 5 h comparisons. $\Delta log_2$ values are calculated by $Log_2$ fold change for Δ*acrB* minus $Log_2$ fold change for the wild type. Positive values indicate more upregulation or less downregulation in wt, negative values more upregulation or less downregulation in *acrB*. Genes are sorted in decreasing $\Delta log_2$ 1vs3 h order.

**ID**: locus name.
**Gene name**
**Operon structure:** if unknown are inferred from genomic sequence and comparison with ecocyc.com.
**STM**: STM gene number.
**Product**: Function of gene product, if known.
**Comments**: General classification of function of gene product.
**Regulation in *Salmonella*:** known regulation, as referenced.
**Regulation in *E. coli***: sourced from Ecocyc.com unless otherwise referenced.
**SalComMac $\log_2$ Anaerobic shock**: Gene expression data from SalComMac (http://bioinf.gen.tcd.ie/cgi-bin/salcom.pl?db=salcom_mac_HL) for the anaerobic shock condition; “Growth in Lennox broth to $OD_{600}$ 0.3 (50 ml), then filled into 50 ml closed Falcon tube and incubated without agitation at 37°C for 30 min.” (8).

**Colour mapping:**

| -4 | -3 | -2 | -1 | 0 | 1 | 2 | 3 | 4 |
|---|---|---|---|---|---|---|---|---|

$\text{Log}_2$ fold change values that are non-significant ($p_{adj} > 0.05$) are in white and marked with a dagger †. $\Delta\text{Log}_2$ fold change values derived from one or more non-significant $\text{Log}_2$ fold change values are also in white.

### Supplemental Table S3. Genes in group 3 - Genes that are more upregulated at 3hr in *wt* than Δ*acrB*

| Log$_2$ fold change | | | | Δ*acrB* – wt Δlog$_2$ | | | | | | | | |
|---|---|---|---|---|---|---|---|---|---|---|---|---|
| wt 1vs3 | wt 3vs5 | acrB 1vs3 | acrB 3vs5 | 1vs3 | 3vs5 | ID | Gene | Operon | STM | Product | Comments | Regulation in *E. coli* |
| 7.28 | -2.01 | 2.33 | 2.80 | -4.95 | 4.81 | SL1344_RS21095 | *metF* | *metF* | STM4105 | Methylenetetrahydrofolate reductase | Methionine / SAM synthesis | MetJ repressed |
| 5.70 | -3.19 | 0.77 | 1.46 | -4.93 | 4.64 | SL1344_RS20380 | *metR* | *metR* | STM3964 | HTH-type transcriptional regulator MetR | Methionine / SAM synthesis | MetJ, MetR repressed |
| 7.88 | -1.84 | 3.10 | 3.65 | -4.78 | 5.49 | SL1344_RS20385 | *metE* | *metE* | STM3965 | 5 - methyltetrahydropteroyltriglutamat e homocysteine S-methyltransferase | Methionine / SAM synthesis | MetJ repressed, MetR, OxyR activated |
| 5.42 | -1.62 | 1.00 | 2.79 | -4.42 | 4.40 | SL1344_RS21470 | *metA* | *metA* | STM4182 | Homoserine O-succinyltransferase | Methionine / SAM synthesis | MetJ repressed |
| 4.79 | -3.08 | 0.44 | 1.50 | -4.35 | 4.58 | SL1344_RS21070 | *metL* | *metBL* | STM4101 | Bifunctional aspartate kinase/homoserine dehydrogenase II | Methionine / SAM synthesis | MetJ repressed, PhoP activated |
| 6.20 | -3.28 | 1.87 | 1.61 | -4.33 | 4.88 | SL1344_RS03045 | *ybdL/ mtnE* | *ybdL/mtnE* | STM0603 | Methionine aminotransferase | Methionine salvage pathway from polyamine biosynthesis pathway. | Lrp repressed<br>MetJ regulated? (9) |
| 4.74 | -3.74 | 0.44 | 0.96 | -4.30 | 4.70 | SL1344_RS21075 | *(mscS)* | | STM4102 | Pseudogene - mechanosensitive ion channel | Pseudogene, contains a stop codon | |
| 5.47 | -5.17 | 1.46 | -0.62 | -4.01 | 4.55 | SL1344_RS19860 | *pstS* | *pstSCAB-phoU* | STM3857 | Phosphate ABC transporter substrate-binding protein PstS | High affinity phosphate uptake | RpoS,FNR,IHF,PhoB activated, Nac repressed |
| 5.91 | -2.37 | 1.91 | 2.02 | -4.00 | 4.39 | SL1344_RS03040 | *ybdH* | *ybdH* | STM0602 | Putative glycerol dehydrogenase | Glycerol utilisation? | MetJ regulated? (9) |
| 4.74 | -3.44 | 1.23 | -0.26 † | -3.51 | 3.18 | SL1344_RS19850 | *pstA* | *pstSCAB-phoU* | STM3855 | Phosphate ABC transporter permease PstA | High affinity phosphate uptake | RpoS,FNR,IHF,PhoB activated, Nac repressed |
| 4.50 | -3.37 | 1.17 | -0.16 † | -3.32 | 3.21 | SL1344_RS19855 | *pstC* | *pstSCAB-phoU* | STM3856 | Phosphate ABC transporter permease PstC | High affinity phosphate uptake | RpoS,FNR,IHF,PhoB activated, Nac repressed |
| 3.65 | -0.12 † | 0.51 | 2.29 | -3.14 | 2.41 | SL1344_RS03035 | *ybdD* | *ybdD* | STM0601 | YbdD/YjiX family protein | Pyruvate import in *E. coli* | |
| 4.12 | -2.17 | 1.02 | 0.76 | -3.10 | 2.92 | SL1344_RS01250 | *metN* | *metNIQ* | STM0247 | Methionine ABC transporter ATP-binding protein MetN | Methionine uptake / SAM synthesis | MetJ repressed, HypT activated |
| 4.00 | -3.50 | 1.16 | -0.52 | -2.85 | 2.98 | SL1344_RS19845 | *pstB* | *pstSCAB-phoU* | STM3854 | Phosphate ABC transporter ATP-binding protein PstB | High affinity phosphate uptake | RpoS,FNR,IHF,PhoB activated, Nac repressed |
| 3.83 | -1.88 | 1.07 | 1.14 | -2.76 | 3.02 | SL1344_RS03050 | *ybdM* | *ybdNM* | STM0604 | Transcriptional regulator related to SpoOJ (10) | Function unknown | |
| 1.62 | -0.95 | -1.01 | 1.20 | -2.63 | 2.15 | SL1344_RS22020 | *proP* | *proP* | STM4290 | Proline / glycine betaine transporter | Osmotic stress response | RpoS, Lrp, Fis activated |
| 3.42 | -1.33 | 0.90 | 0.85 | -2.52 | 2.18 | SL1344_RS16345 | *metC* | *metC* | STM3161 | Cystathionine beta-lyase | Methionine / SAM synthesis | MetJ repressed |
| 3.59 | -1.43 | 1.26 | 1.02 | -2.32 | 2.45 | SL1344_RS01245 | *metI* | *metNIQ* | STM0246 | Methionine ABC transporter permease MetI | Methionine uptake / SAM synthesis | MetJ repressed, HypT activated |
| 3.47 | -0.74 † | 1.26 | 1.08 | -2.21 | 1.83 | SL1344_RS21100 | *katG* | *katG* | STM4106 | Catalase/peroxidase HPI | ROS defence - catalase | Induced on entry into stationary phase (11) |
| 2.49 | -0.26 † | 0.48 | 1.83 | -2.02 | 2.10 | SL1344_RS21475 | *aceB* | *aceBAK* | STM4183 | Malate synthase A | Glyoxylate shunt | Lrp, Cra, IHF activated, ArcA, Nac, CRP, IclR repressed |
| 1.11 | -0.92 | -0.73 | 1.08 | -1.84 | 2.00 | SL1344_RS04365 | *yliJ / gstB* | *yliJ* | STM0862 | Putative glutathione S-transferase | Dehalogenates Br-acetate and I-acetate in *E. coli* | |
| 3.27 | -0.87 † | 1.45 | 0.57 † | -1.82 | 1.44 | SL1344_RS25905 | *yqgD* | *yqgD* | STM3089 | Hypothetical protein | Function unknown | |

| | | | | | | | | | | | | |
|---|---|---|---|---|---|---|---|---|---|---|---|---|
| 1.24 | 0.92 † | -0.50 | 2.05 | -1.74 | 1.13 | SL1344_RS05680 | *msyB* | *msyB* | STM1153 | SecY/SecA suppressor protein | Periplasmic protein export | RpoS activated (12) |
| 0.98 | 0.35 † | -0.69 | 1.59 | -1.67 | 1.24 | SL1344_RS21380 | *rsd* | *rsd* | STM4165 | Anti-RNA polymerase sigma 70 factor | Coordinates entry to stationary phase. | Lrp,McbR,RcdA,SdiA,SlyA activated, ArcA Nac repressed |
| -0.83 | 0.00 † | -2.45 | 0.00 † | -1.62 | 0.00 | SL1344_RS14535 | *nrdI* | *nrdHI* | STM2806 | Ribonucleotide reductase assembly protein NrdI | Nucelotide metabolism | IscR activated, Fur and NrdR repressed |
| 1.10 | 2.36 | -0.47 | 2.99 | -1.58 | 0.63 | SL1344_RS01875 | *yahO* | *yahO* | STM0366 | Hypothetical protein | Function unknown | RpoS, ppGpp activated |
| 3.46 | -0.81 † | 1.89 | 0.99 | -1.57 | 1.80 | SL1344_RS15985 | *metK* | *metK* | STM3090 | Methionine adenosyltransferase | SAM synthesis | MetJ repressed, CRP repressed |
| 1.03 | 0.64 † | -0.53 | 1.46 | -1.56 | 0.82 | SL1344_RS05675 | *yceK* | *yceK* | STM1152 | Hypothetical OM lipoprotein | LPS assembly in *E. coli* (13) | RpoS activated (12) |
| 1.89 | -0.93 † | 0.43 | 1.13 | -1.46 | 2.06 | SL1344_RS02595 | *sfbA* | *sfbABC* | STM0510 | Metal ABC transporter substrate-binding protein | Iron uptake? | |
| 1.80 | 0.13 † | 0.35 | 2.03 | -1.45 | 1.90 | SL1344_RS11210 | *yohK* | *yohJK* | STM2182 | Putative 3-hydroxypropanoate export protein | Hydroxypropanoate export? | |
| 0.93 | 0.62 † | -0.45 | 1.55 | -1.38 | 0.93 | SL1344_RS10650 | *yeeZ* | *yeeZ* | STM2070 | NAD(P)-dependent oxidoreductase | Unknown function | |
| 0.95 | -0.12 † | -0.42 | 0.98 | -1.37 | 1.09 | SL1344_RS00815 | *yacL* | *yacL* | STM0160 | Protein YacL | Unknown function | Lrp activated |
| 0.91 | 0.01 † | -0.40 | 1.41 | -1.31 | 1.41 | SL1344_RS04590 | *clpA* | *clpSA* | STM0945 | ATP-dependent Clp protease ATP-binding subunit | Proteolysis | |
| 1.68 | -0.62 † | 0.39 | 0.47 † | -1.29 | 1.10 | SL1344_RS11205 | *yohJ* | *yohJK* | STM2181 | Putative 3-hydroxypropanoate export protein | Hydroxypropanoate export? | |
| 1.60 | 0.18 † | 0.34 | 1.34 | -1.26 | 1.16 | SL1344_RS06585 | *ghoS* | *ghoS* | STM1326 | Type V toxin-antitoxin system endoribonuclease antitoxin GhoS | TA system | |
| 2.30 | 3.74 | 1.05 | 4.61 | -1.25 | 0.87 | SL1344_RS05235 | *rmf* | *rmf* | STM1066 | Ribosome modulation factor | Converts ribosomes to dimeric form in stationary phase | ppGpp activated, ArcA repressed |
| 1.70 | 1.86 | 0.47 | 2.41 | -1.23 | 0.55 | SL1344_RS06940 | *sseA* | SPI-2 | STM1397 | SPI-2 type III secretion system chaperone SseA | SPI-2 | |
| 0.77 | -1.13 | -0.46 | -0.02 † | -1.23 | 1.11 | SL1344_RS15870 | *yggB / mscS* | *yggB* | STM3067 | small-conductance mechanosensitive channel MscS | Mechanosensing | Lrp repressed |
| 1.61 | 1.69 | 0.40 | 2.19 | -1.21 | 0.50 | SL1344_RS17230 | *yhcH / nanQ* | *nanTEKQ* | STM3335 | N-acetylneuraminate anomerase | Sialic acid metabolism | CRP activated, Fis, NanR repressed |
| 2.30 | 0.69 | 1.09 | 2.01 | -1.20 | 1.32 | SL1344_RS18435 | *tcp* | *tcp* | STM3577 | Methyl-accepting chemotaxis protein II | Chemotaxis | |
| 2.89 | -1.36 | 1.69 | -0.52 | -1.20 | 0.84 | SL1344_RS16195 | *hcp* | *hcp-hcr* | STM3131 | Type VI secretion system tube protein Hcp | Protein secretion | |
| 0.81 | 0.58 † | -0.36 | 1.17 | -1.17 | 0.59 | SL1344_RS06900 | *orf319* | SPI-2 | STM1389 | hypothetical protein | SPI-2 | |
| 1.51 | -1.05 | 0.34 | 0.49 | -1.17 | 1.54 | SL1344_RS18795 | *ghrB* | *ghrB* | STM3646 | Glyoxylate / hydroxypyruvate reductase | Gluconate metabolism | |
| 1.65 | -0.31 † | 0.51 | 0.91 | -1.14 | 1.21 | SL1344_RS11875 | *cheV* | *cheV* | STM2314 | Chemotaxis protein CheV | Chemotaxis | |
| 1.58 | 1.28 | 0.45 | 1.70 | -1.13 | 0.43 | SL1344_RS06995 | *ssaI* | SPI-2 | STM1408 | EscI/YscI/HrpB family type III secretion system inner rod protein | SPI-2 | |
| 0.85 | -0.33 † | -0.27 | 1.16 | -1.12 | 1.48 | SL1344_RS04585 | *clpS* | *clpSA* | STM0944 | ATP-dependent Clp protease adapter ClpS | Proteolysis | Lrp activated, PhoP repressed |
| 0.69 | 0.19 † | -0.41 | 1.17 | -1.09 | 0.98 | SL1344_RS19370 | *cigR* | *cigR* | STM3762 | Anti-virulence regulatory inner membrane protein | Regulator of virulence | |
| 0.73 | 0.72 † | -0.35 | 0.95 | -1.07 | 0.23 | SL1344_RS06575 | *yniB* | *yniB* | STM1323 | Hypothetical protein | Unknown function | |
| 1.59 | 0.29 † | 0.52 | 1.18 | -1.07 | 0.88 | SL1344_RS02190 | *phnT* | *phnSTUV* | STM0428 | 2-aminoethylphosphonate ABC transport system ATP-binding subunit PhnT | Aminoethylphosphonate import | |
| 2.24 | -0.31 † | 1.18 | 0.63 | -1.06 | 0.93 | SL1344_RS18505 | *uspA* | *uspA* | STM3591 | Universal stress protein UspA | Stress response | IHF, ppGpp activated, Nac, FadR repressed |

| | | | | | | | | | | | |
|---|---|---|---|---|---|---|---|---|---|---|---|
| 1.60 | 1.25 | 0.54 | 1.55 | -1.06 | 0.30 | SL1344_RS06990 | *ssaH* | SPI-2 | | EscG/YscG/SsaH family type III secretion system needle protein co-chaperone | SPI-2, HilA repressed (14) |
| 0.83 | -0.04 † | -0.23 | 0.58 | -1.06 | 0.62 | SL1344_RS12340 | | RS12340 | STM2404 | Ion channel protein | |
| 0.57 | 0.56 | -0.43 | 0.89 | -1.00 | 0.33 | SL1344_RS18425 | *yhhN* | *yhhN* | STM3575 | Hypothetical protein | Unknown function |

**Key to columns (left to right)**

**$Log_2$ fold change** values for each gene: 1 h vs 3 h in the wild type; 3 h vs 5 h in the wild type; 1 h vs 3 h in Δ*acrB*; 3 h vs 5 h in Δ*acrB*.

**$\Delta Log_2$ values** for the 1 h vs 3 h and 3 h vs 5 h comparisons. $\Delta log_2$ values are calculated by $Log_2$ fold change for Δ*acrB* minus $Log_2$ fold change for the wild type. Positive values indicate more upregulation or less downregulation in wt, negative values more upregulation or less downregulation in *acrB*. Genes are sorted in decreasing $\Delta log_2$ 1vs3 h order.

**ID**: locus name.

**Gene name**

**Operon structure:** if unknown are inferred from genomic sequence and comparison with ecocyc.com.

**STM**: STM gene number.

**Product**: Function of gene product, if known.

**Comments**: General classification of function of gene product.

**Regulation in *E. coli***: sourced from Ecocyc.com unless otherwise referenced.

**Colour mapping:**

| -4 | -3 | -2 | -1 | 0 | 1 | 2 | 3 | 4 |
|---|---|---|---|---|---|---|---|---|

$Log_2$ fold change values that are non-significant ($p_{adj} > 0.05$) are in white and marked with a dagger †. $\Delta Log_2$ fold change values derived from one or more non-significant $Log_2$ fold change values are also in white.
